# Anteromedial Thalamus Gates the Selection & Stabilization of Long-Term Memories

**DOI:** 10.1101/2023.01.27.525908

**Authors:** Andrew C. Toader, Josue M. Regalado, Yan Ran Li, Andrea Terceros, Nakul Yadav, Suraj Kumar, Sloane Satow, Florian Hollunder, Alessandra Bonito-Oliva, Priya Rajasethupathy

## Abstract

Memories initially formed in hippocampus gradually stabilize to cortex, over weeks-to-months, for long-term storage. The mechanistic details of this brain re-organization process remain poorly understood. In this study, we developed a virtual-reality based behavioral task and observed neural activity patterns associated with memory reorganization and stabilization over weeks-long timescales. Initial photometry recordings in circuits that link hippocampus and cortex revealed a unique and prominent neural correlate of memory in anterior thalamus that emerged in training and persisted for several weeks. Inhibition of the anteromedial thalamus-to-anterior cingulate cortex projections during training resulted in substantial memory consolidation deficits, and gain amplification more strikingly, was sufficient to enhance consolidation of otherwise unconsolidated memories. To provide mechanistic insights, we developed a new behavioral task where mice form two memories, of which only the more salient memory is consolidated, and also a technology for simultaneous and longitudinal cellular resolution imaging of hippocampus, thalamus, and cortex throughout the consolidation window. We found that whereas hippocampus equally encodes multiple memories, the anteromedial thalamus forms preferential tuning to salient memories, and establishes inter-regional correlations with cortex, that are critical for synchronizing and stabilizing cortical representations at remote time. Indeed, inhibition of this thalamo-cortical circuit while imaging in cortex reveals loss of contextual tuning and ensemble synchrony in anterior cingulate, together with behavioral deficits in remote memory retrieval. We thus identify a thalamo-cortical circuit that gates memory consolidation and propose a mechanism suitable for the selection and stabilization of hippocampal memories into longer term cortical storage.

## INTRODUCTION

The hippocampus has a well-established role in the initial formation and storage of memory (Kandel et al., 2014). However, little is understood about brain mechanisms that support the selection and re-organization of memories into longer-term storage. Early discoveries demonstrated that the hippocampus may have a time-limited role in memory (Scoville and Milner, 1957; Zoia-Morgan and Squire, 1990; Kim and Fanselow, 1992) and that the cortex may be the permanent repository of long-term memories (Squire and Alvarez, 1995; McClelland et al., 1995; Nadel and Moscovitch, 1997; Dudai, 2004; Frankland and Bontempi, 2005). To formalize how this memory re-organization may occur, a standard model of systems consolidation was proposed, in which the hippocampus transiently stores new memories, and over time, trains cortex, in particular anterior cingulate cortex, to create and store more enduring representations (Buzsáki, 1996; Siapas and Wilson, 1998; Trachtenberg et al., 2002; Maviel at al., 2004; Frankland et al., 2004; Wiltgen et. al, 2004; Sacco and Sacchetti, 2010). Subsequent studies led to alternative models, which proposed that memories are initially created in both hippocampus and cortex (Tse et al., 2007; Tse et al., 2011; Lesburguères et al., 2011; Bero at al., 2014, Kitamura et al. 2017) (though the nature of each representation may differ) (Nadel and Moscovitch, 1997; Yonelinas et al., 2019; Barry and Maguire, 2019), and over time, hippocampal representations are either progressively reduced or continuously maintained together with cortical representations. Distinguishing or optimally integrating these models requires detailed study of inter-connected circuits between hippocampus and cortex, that may select, gate, and transfer memory into long term storage throughout the consolidation process; an area that has lacked investigation and mechanistic insight. One major challenge is the technical limitations preventing real-time exploration of dynamically evolving memory ensembles in hippocampus, cortex, and intervening circuits simultaneously and longitudinally in the behaving animal. Here, we developed a memory-guided behavioral task that mice perform reliably in the head-fixed setting for months, and performed anatomical tracing, neural dynamic recordings, and optogenetic perturbations to characterize the role of these intervening circuits in memory consolidation. We then developed a method to perform cellular resolution imaging across multiple functionally connected but neuro- anatomically distributed circuits, over months-long timescales, to contribute new insights into how memories are selected and stabilized into long-term storage.

## RESULTS

### A virtual-reality-based memory consolidation behavioral task

We began by designing a contextual memory-guided task that mice can perform in a head-fixed setting to enable the precise tracking of stimuli and behavior during high resolution brain imaging. The majority of memory consolidation studies thus far have used fear conditioning as a standard behavioral model due to its robustness and ease of implementation. However, since important differences may exist between the consolidation of implicit versus explicit memories (Kandel et al. 2014), we designed a more explicit task, by requiring task engagement through self-initiation of trials (Figure 1) and by requiring multi-modal contextual integration (Figure S1F) and spatial navigation (Figures S2A-B) for successful task performance. In brief, mice navigated on an axially-fixed track ball in a virtual-reality environment (Harvey et al., 2009; Rajasethupathy et al., 2015) composed of a corridor with three distinct zones: start zone, cue zone, and outcome zone (Figures 1A and 1B). Mice initiated trials by entering the cue zone where they received multi-modal contextual cues signifying one of two potential contexts, where one predicted sucrose reward while the other predicted aversive airpuff in the outcome zone. Mice underwent a training phase over five days and were subsequently probed with weekly retrieval sessions throughout the consolidation phase (Figure 1B). On each day, once a session was initiated, the task was entirely automated requiring no human intervention. By employing appropriate shaping procedures (Figure S1A, Methods), we observed high task engagement, learning, and memory that persisted for weeks to months (Figures 1C-E). While in the cue zone, animals demonstrated learning and memory by reliably decreasing their speed and anticipatory licking when cues predicted aversive airpuff, while increasing speed and anticipatory lick rate when cues predicted sucrose reward (Figures 1C and S1B). On average, mice demonstrated a high degree of learning and memory retrieval using both measures quantified as differences in absolute values or normalized as a discrimination index (DI, Methods) (Figures 1D and S1C-E; 1.7 vs 0.7 Z-scored speed in reward vs aversive context on day 55, P<0.001, two-way ANOVA with repeated measures; 2.5Hz vs 0.2 Hz lick rate in reward vs aversive context on day 55, P<0.01, two-way ANOVA with repeated measures). Importantly, neither sets of cues, in the absence of paired reinforcement, were inherently rewarding or aversive (Figure 1C, Day 0). To minimize extinction due to repeated longitudinal retrieval sessions, we performed minimal retraining (Figure 1C, Methods), which we confirmed yields comparable performance to a single remote retrieval session (Figure 1E), without loss of motivation when reward is again provided (Figure S1D).

**Figure 1.**
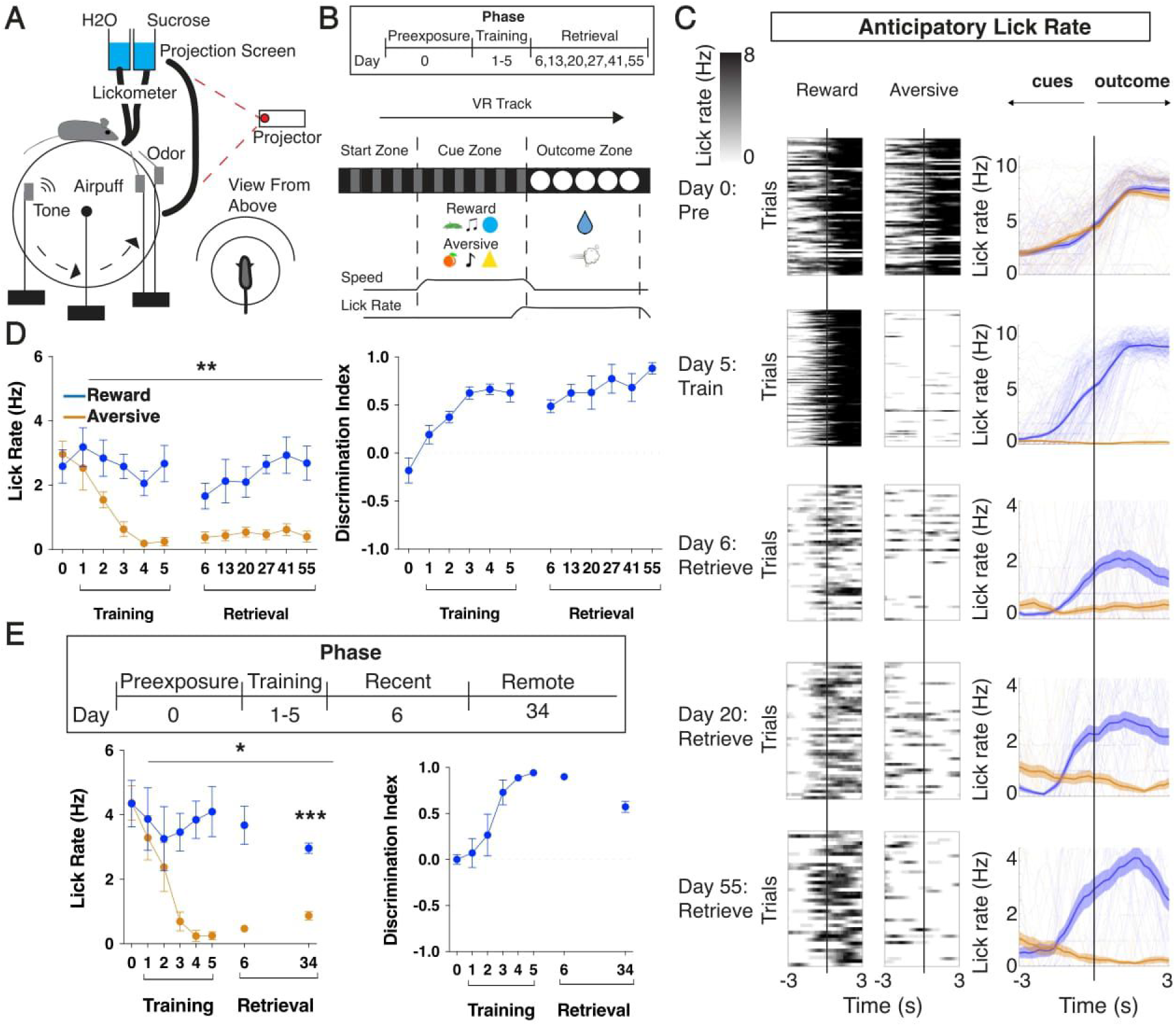
A virtual-reality based contextual memory task spanning months. (A) Schematic of virtual reality experimental setup. (B) Timeline of behavioral task from preexposure (day 0), through training (days 1-5), and retrieval (probe trials on days 6, 13, 20, 27, 41, and 55). A single retraining session occurs after each retrieval session. Virtual reality linear track with start, cue, and outcome zones, and example behavioral parameters (speed, lick rate) tracked. Reinforcements are provided in the outcome zone during training but omitted during retrieval probes. (C) Raster of individual trials and plot of trial averages. Anticipatory lick rate (Hz) is described as the lick rate prior to the outcome zone. Also shown is lick rate 3 seconds into the outcome zone. N=5 mice, 8 trials/context/­ session during training, 4 trials/context/session during retrieval. (D) Quantification of average anticipatory lick rate (Hz) 2 seconds prior to outcome zone entry in each context during preexposure, training, and weekly retrieval sessions. Quantification of discrimination index (difference of anticipatory lick rates in reward and aversive context divided by their sum). N=5 mice, **p<0.01, Two way ANOVA with repeated measures, data are mean± s.e.m. (E) Same as (D), but without weekly retrieval sessions (see also Fig. S6), and testing only recent (day 6) and remote (day 34) retrieval, N=4 mice, *p<0.05, Two way ANOVA with repeated measures, data are mean± s.e.m. ***p<0.001 for R34, Bonferroni corrected for multiple comparisons, data are mean± s.e.m.

To test whether the hippocampus is required to learn and retrieve contextual memories in this task, we expressed the inhibitory opsin stGtACR2 (Mahn et al., 2018) bilaterally in CA1 and delivered light during the cue and outcome zones throughout training (Figure 2A, Figure S2C-D). Hippocampal inhibition during training prevented learning of context discrimination when compared to controls (Figure 2B, see also Figure S2E-F). Furthermore, when mice were trained on the task without inhibition and then subsequently tested the next day (Figure 2C), online hippocampal inhibition disrupted recall (Figure 2D, see also Figure S2G-H). We next tested whether memory in this task undergoes cortical consolidation. To do so, we prepared new cohorts of mice for training, and after three weeks, tested remote recall while performing chemogenetic inhibition (Armbruster et al. 2007) in hippocampus or anterior cingulate cortex (ACC) (Figure 2E). We found that successful recall of remote memories no longer required the hippocampus, but rather critically depended on ACC (Figure 2F). Thus, we established a head-fixed self-initiated contextual memory task that is 1) dependent on the hippocampus for acquisition and recent retrieval, 2) becomes independent of the hippocampus over weeks and cortically dependent, and 3) enables the reliable tracking of memory longitudinally for many weeks.

**Figure 2.**
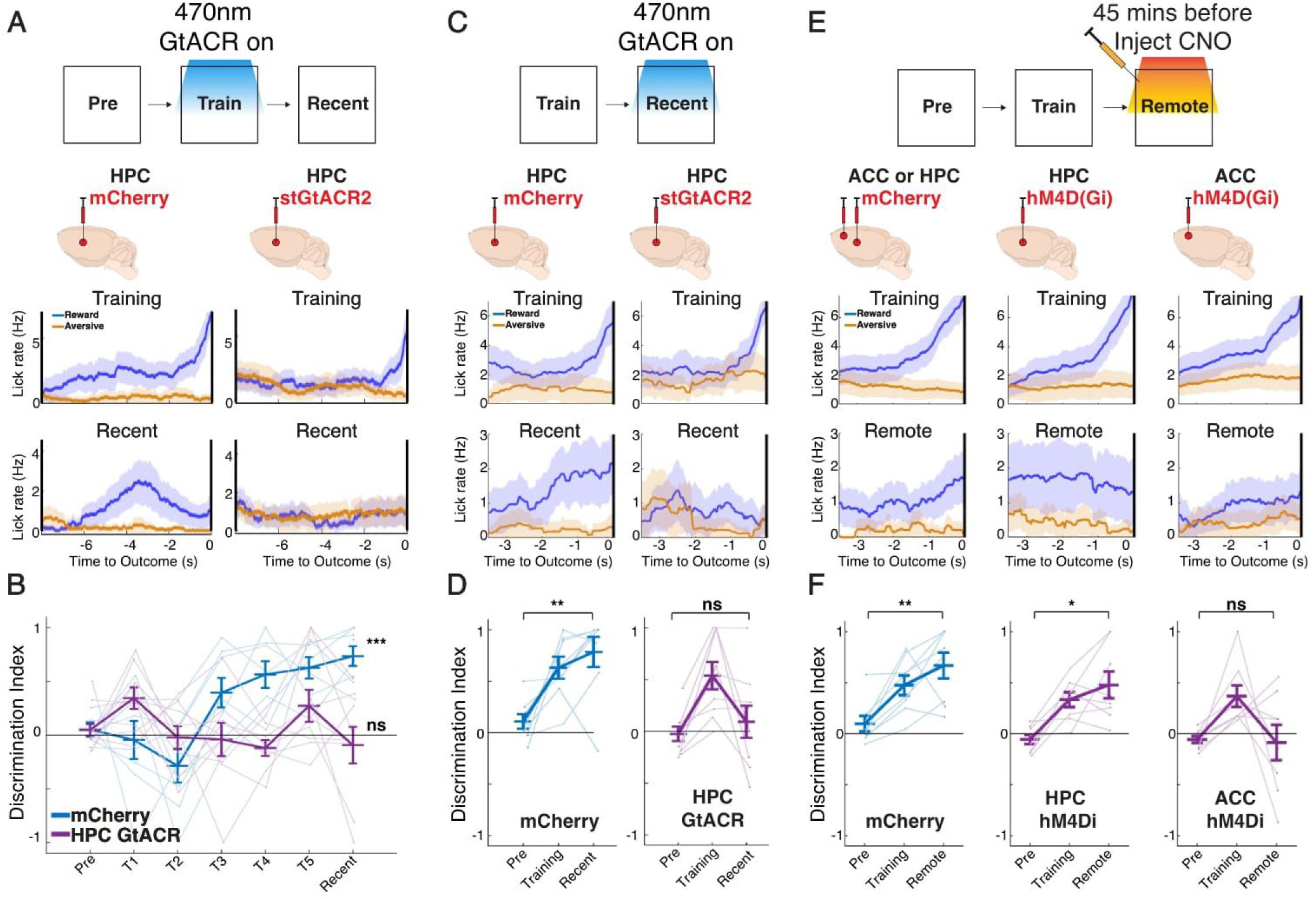
Learning and recent memory recall require HPC but remote recall becomes ACC dependent. (A) Schematic of experimental design: stGtACR2-based optogenetic inhibition during training (T1-T5), followed by a test of recent memory. Raw lick traces are shown for mCherry (HPC with no opsin control, N=8) and GtACR (HPC with opsin, N=8) in each context during training and recent retrieval sessions, data are mean (solid line)± s.e.m (shaded area). (B) Quantification of discrimination between reward and aversive lick rates per mouse cohort on preexposure, training, and recent retrieval sessions *** p<0.001 between pre-exposure and recent for mCherry vs p=0.9780 for HPC GtACR, one-way repeated measures ANOVA with Tukey’s multiple comparison test. Individual data points shown, with mean± s.e.m. (C) Schematic of experimental design: GtACR-based optogenetic inhibition during recent memory in animals that were trained without optogenetic inhibition. Raw lick traces are shown for mCherry (HPC with no opsin control, N=8) and GtACR (HPC with opsin, N=8) in each context during training and recent retrieval sessions, data are mean (solid line)± s.e.m (shaded area). (D) Quantification of discrimination between reward and aversive lick rates per mouse cohort on preexposure, training, and recent retrieval sessions ** p<0.01 for between pre-exposure and recent for mCherry vs p=0.5812 for HPC GtACR, one-way repeated measures ANOVA with post-hoc Tukey’s multiple comparison test. Individual data points shown, with mean± s.e.m. (E) Schematic of experimental design: DREADDS-based chemogenetic inhibition during remote memory in trained animals. CNO was adminis­ tered 45 minutes prior to the remote retrieval session. Raw lick traces are shown for mCherry (ACC or HPC with no hM4Di, N=8), HPC hM4Di (HPC with hM4Di, N=8), ACC hM4Di (ACC with hM4Di, N=8) in each context during training and recent retrieval sessions, data are mean (solid line)± s.e.m (shaded area). (F) Quantification of discrimination between reward and aversive lick rates per mouse cohort on preexposure, training, and remote retrieval sessions. **p<0.01 for mCherry between preexposure and remote, *p<0.05 for HPC hM4Di, p=0.945 for ACC hM4Di, one-way repeated measure ANOVA with post-hoc Tukey’s multiple comparison test. Individual data points shown, with mean± s.e.m.

### Longitudinally tracking neural correlates of memory consolidation

To select regions of interest to monitor via neural recordings during memory consolidation, we began by identifying brain regions which send direct projections to anterior cingulate cortex. Brain-wide retrograde tracing from anterior cingulate cortex using retroAAV revealed many inputs, of which we focused on regions that in turn have known inputs from hippocampus (HPC), including anterior thalamus (ANT, via mammillary bodies), basolateral amygdala (BLA), and entorhinal cortex (ENT) (Maren, 2001; Buzsáki and Moser, 2013; Jankowski et al. 2013) (Figure 3A and S3A). We injected the genetically encoded calcium indicator GCaMP6f in ACC, HPC, ANT, BLA, and ENT, implanted optical fibers above each region, and recorded neural activity during the task daily (during training) or weekly (during consolidation) by multi-fiber photometry (Kim et al., 2016; Sych et al., 2019). The injection coordinates (Methods) were optimized to target the area of ANT, BLA, and ENT that contained direct projections to ACC. Neural activity recordings (10 Hz) from each brain region of a given animal were then frame projected onto a camera sensor, and custom MATLAB scripts (Methods) were used to extract time-series data, regress out motion-related artifacts and align to behavioral data at 100-ms resolution (Figure 3B, Methods). To ensure high-quality data, we characterized signal-to-noise (Figure S3B) and excluded recordings below a statistically defined threshold (Figures S3C-D; Methods).

**Figure 3.**
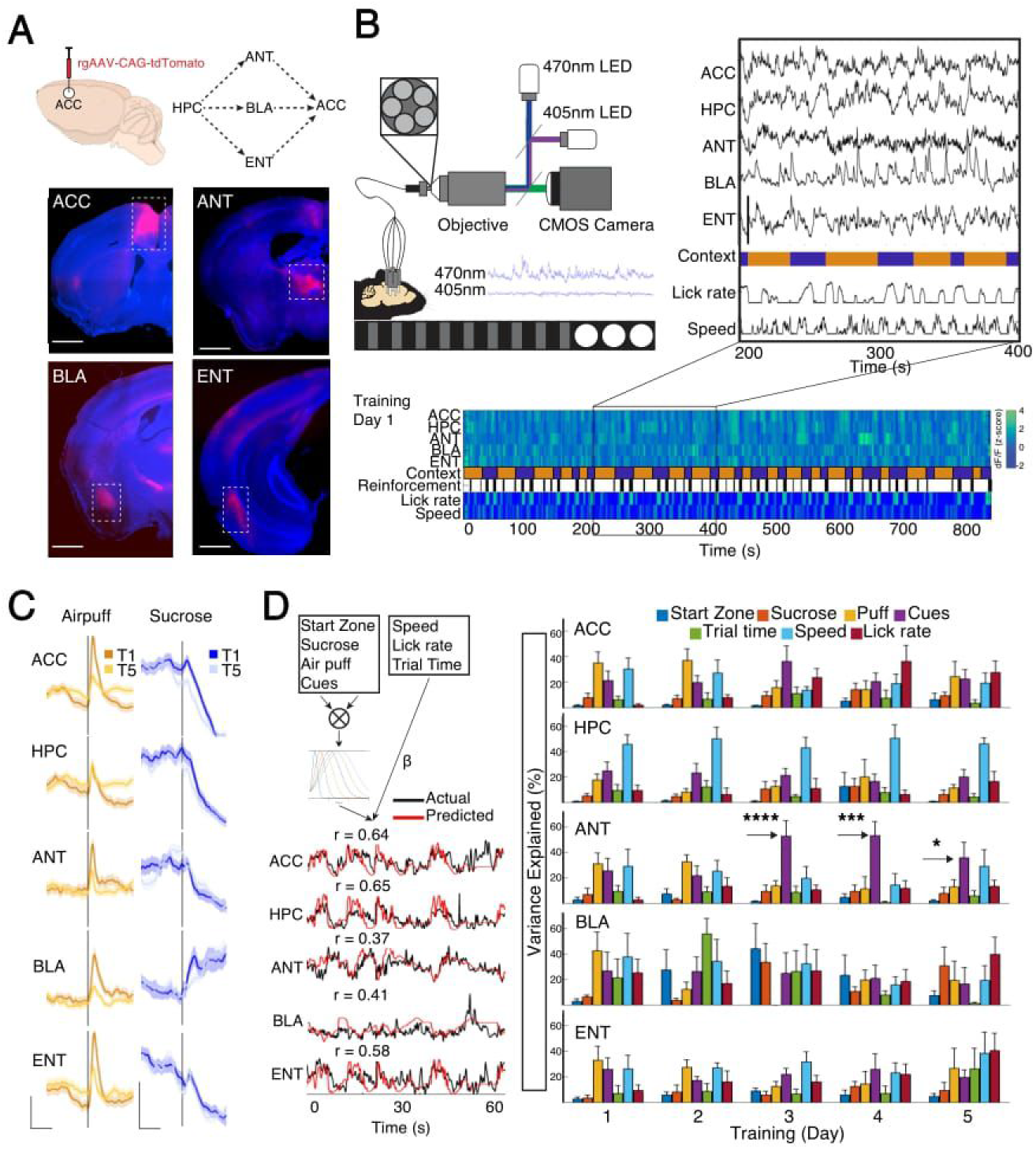
Longitudinal photometry recordings with distributed and mixed coding of task-relevant features. (A) rgAAV CAG tdTomato was injected into ACC, revealing retrogradely labeled neurons in anterior thalamus (ANT), basolateral amygdala (BLA), and entorhinal cortex (ENT) among other regions. Each of ANT (via mammillary bodies), BLA, and ENT receive input from the dorsal hippo­ campus (HPC). Scale: 1 mm. (B) Left: Photometry setup Excitation light is delivered through fiber optic cables placed above the five implanted cannulas, and emitted fluorescence is projected back onto a CMOS camera. The calcium dependent fluores cence from GCaMP6f is normalized by the calcium independent fuores­ cence to correct for movement artifacts and calculate △F/F Bottom and inset: Example traces of △F/F from one mouse on training day 1 from all five regions aligned to task and behavioral variables (C) Robust trial-averaged responses to reinforcement (air puff or sucrose) on training day 1 (T1) and 5 (T5). N=7 mice. Data are mean (dark line) and s e m. (shaded area). Photometry scale: x/y: 2s/1z (D) Left: Schematic of the generalized lnear used to predict bulk neural activity from task and behavioral varables, w^i^th examp e mode outputs from each region shown below (see methods). Right: Percentage of varance explaned in the mode by each variable throughout training (T1-T5) for each region N=7 mice Arrows: One way ANOVA w ^i^th Tukey’s multiple comparison test between inter tria interval (start zone) and cue zone for T3 (****p<0 0001), T4 (***p<0.001) and T5 (*p<O.0S). Data are mean ± s.e.m. **V**

We first asked whether we could identify neural signals in these regions that were tuned to task-relevant features. We observed several types of neural responses including those that were time locked to onset of reinforcement, including air puff and sucrose (Figure 3C). Given that most brain regions encoded multiple variables simultaneously, we developed an encoding model, a generalized linear model, to isolate signals related to each measured task variable, including task zone, sensory cues, reinforcement, elapsed time, as well as the continuous behavioral variables including run speed and lick rate. The model reliably predicted neural activity time series on held-out trials across regions and across days (Figures 3D and S4A). Using this model, we calculated the variance explained in the predicted activity by each task variable (Methods). We observed that multi-modal information existed in all regions, some of which were modulated by learning and time. For instance, air puff representations diminished in all five regions across days, while sucrose representations increased in BLA (Figure 3D). Notably, a significant cue representation, relative to the start zone inter-trial interval, and independent of running or licking, uniquely emerged in ANT on days 3-5 of training (Figure 3D, arrows). Further characterization revealed that these changes in task-related tuning were not due to changes in the overall magnitude or rate of neural population activity, but rather changes in the relative timing to contextual and behavioral variables. (Figures S4B-D).

We proceeded to identify the emergence and evolution of “memory-related” neural activity patterns by focusing on neural activity that tracks both the conditioned stimulus, i.e., cues, and conditioned response, i.e., anticipatory lick rate. We found that neural activity in ANT contained prominent signals for both (Figure 4). Specifically, ANT contained cue-related signals that were distinct for each context, which emerged in training and persisted through consolidation, and which were not explained by changes in animal speed (Figures 4A and 4B). Importantly, these signals emerged late in training, ~T4-T5, demonstrating that it is not a sensory signal, but rather a representation of the learned association. Strikingly, ANT also contained a prominent neural activity signal tracking the conditioned response, i.e., anticipatory lick, when aligned to outcome zone that persisted through consolidation (Figures 4C and 4D). Again, of note, these neural signals emerged on later sessions than the learned anticipatory lick behavior (Figure S4E), and within the cue zone preceded the onset of licking (Figure 4C), which together support that it represents the cognitive process of memory retrieval rather than the motor process of licking. Such memory related signals tracking the conditioned stimulus and conditioned response were not observed in BLA and ENT. In summary, these findings highlight two neural signatures in ANT that together represent a learned association, which is cognitive in nature, rather than sensory or motor. We do not dismiss functional roles of BLA and ENT in memory consolidation because bulk neural activity recordings often obscure fine scale computations; however, given the unique and prominent signals identified in ANT, we proceeded to further characterize its unknown contributions during memory consolidation through causal manipulations and cellular resolution neural activity recordings.

**Figure 4.**
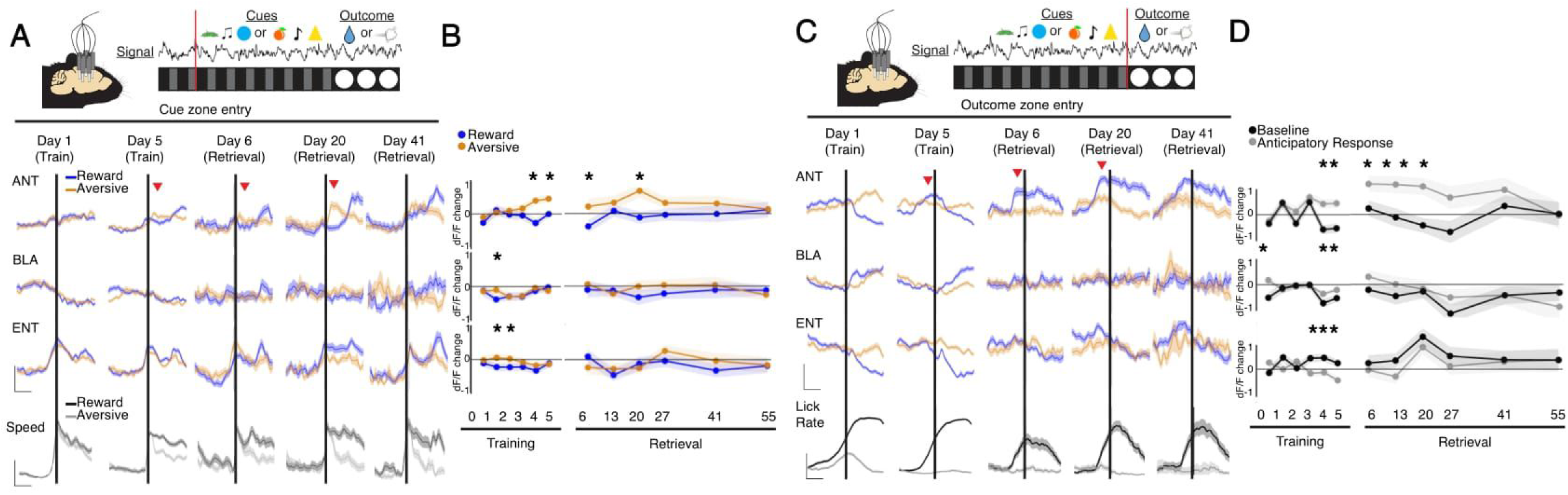
A neural correlate of memory in anterior thalamus persists for weeks. (A) Top: schematic of VR track with cue zone entry in red. Bottom: Mean dF/F in ANT, BLA and ENT aligned to cue zone entry for both contexts (blue vs orange). Bottom row: animal speed aligned to cue zone entry. Red arrow signifies AM signals quantified in 4B (see Methods). N=7 mice. Data are mean (dark line) with s.e.m. (shaded area). Photometry scale: x/y: 2s/1 z. Speed scale: x/y: 2s/0.5z. (B) Quantification of mean change in dF/F at 2s vs. Os in cue zone, assessed separately for each context. N=7 mice, *p<0.05, paired t-test. (C) Same as (A), but aligned to outcome zone entry. Photometry scale: x/y: 2s/1 z. Lick rate scale: x/y: 2s/5Hz. Red arrow signifies AM signals quantified in 4D (see Methods). (D) Quantification of the mean difference in dF/F between contexts prior to outcome zone entry (anticipatory) compared to cue zone entry (baseline). N=7 mice, *p<0.05, paired t-test.

### AM-ACC activity manipulations drive bi-directional changes in memory consolidation

Within the anterior thalamus, we conjectured that the anteromedial (AM) thalamus is anatomically positioned to function in consolidation because it receives direct inputs from the dorsal hippocampus (Wright et al. 2010), and indirect inputs via the mammillary bodies (i.e., Papez circuit), in turn sending direct projections to anterior cingulate cortex (Aggleton et al., 1999, Janowski et al., 2013) (which we confirm via trans-synaptic viral tracing, Figure S3A). To assess whether the AM thalamus is required for memory consolidation, we performed projection-defined optogenetic inhibition of the AM-to-ACC circuit during training. We expressed a soma-targeted inhibitory opsin stGtACR2 by injecting retroAAV- Cre in ACC and floxed-stGtACR2 (or floxed-mCherry for control) in AM thalamus and implanted optical fibers bilaterally above AM thalamus. Importantly, at this dorsal location, no other anterior thalamic nuclei project to ACC, and thus we can confirm an AM-to-ACC specific manipulation. Expression and accurate implantation at the injection site were verified by histology (Figure S5A). We observed that bilateral inhibition of AM-to-ACC during training (Figures 5A) resulted in no significant learning or recent memory deficit, but a significant deficit in remote memory retrieval (Figures 5B-D). This was not due to significant changes in how the mice valued the reward or were motivated to lick, because in a subsequent re-training session following remote memory retrieval, the lick rates to reward in the GTACR cohort matched that of the control cohort (Figure S5C). To ensure our results were not confounded by the intervening recent retrieval with re-training, we repeated this experiment while only testing the mice at remote time (Figure S6A). We again found that mice receiving AM inhibition during training exhibited no learning impairment but substantial impairment of memory recall at remote timepoints (Figures S6B-D). Beyond the training phase, we observed that AM-to-ACC activity was also required at various times during the consolidation window (Figures S6E-H), but due to the repeated nature of the re-trainings and retrievals in this experiment, it is difficult to interpret whether these effects are due to consolidation, extinction or online retrieval. Thus we focused for the remainder of the study on the effects of AM inhibition during training, which while not needed for recent recall demonstrates a clear deficit in memory consolidation and remote recall.

**Figure 5.**
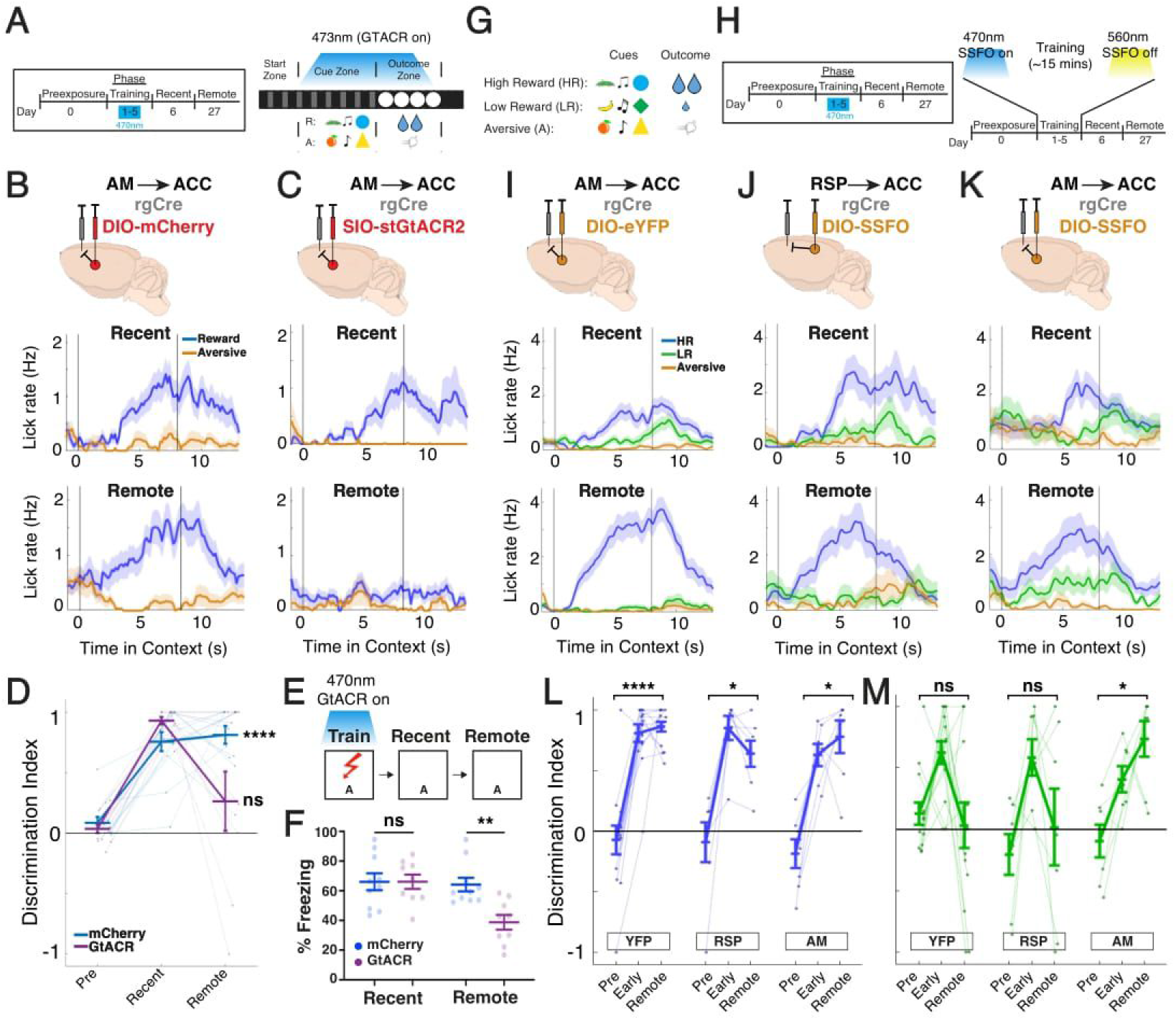
Inhibiting or enhancing AM->ACC activity drives bi-directional changes to remote memory retrieval. **(A)** Schematic of experimental design: stGtACR2-based optogenetic inhibition during training (T1-T5), followed by a test of recent (R6) and remote (R27) memory. Light was delivered during cue and outcome periods of the trial. See also Fig. S6 for optogenetic inhibition during training followed by a direct test of remote memory. (B,C) Injection strategy for targeting anteromedial thalamus (AM) projections to ACC (AM->ACC) in mCherry control and stGtACR2 opsin cohorts. Raw lick traces are shown for each mouse in each context on recent and remote retrieval sessions, data are mean (solid line)± s.e.m (shaded area). (D) Quantification of discrimination between reward and aversive lick rates per mouse on preexpo­ sure, recent, and remote retrieval sessions; mCherry (AM->ACC no opsin control, N=13), GtACR (AM->ACC with opsin, N=9). **’*p<0.0001 for mCherry between Preexposure and Remote, p>0.05 for GtACR, one-way repeated measures ANOVA with post-hoc Tukey’s multiple comparison test. Individual data points shown, with mean± s.e.m. (E) Schematic of experimental design: stGtACR2-based optogenetic inhibition during training, followed by a test of recent and remote memory in a contextual fear conditioning task. (F) Quantification of freezing behavior in training and remote retrieval sessions in mCherry (AM->ACC no opsin control, N=11) and GtACR (AM->ACC with opsin, N=9). **p<0.01 between mCherry and GtACR cohorts on remote, unpaired t-test. Individual data points shown, with mean± s.e.m. (G) Extension of current behavioral task to include two reward contexts (high reward, HR; low reward, LR; aversive, A). (H) Schematic of experimental design: SSFO-based enhancement of neural excitability during training (T1-T5), followed by testing of recent and remote memory. 4 70nm light was delivered at the start of each training session to activate the SSFO and then 560nm light delivered at the end to deactivate the SSFO. See also Fig. S6 for optogenetic activation during training followed by a direct test of remote memory. (1-K) Injection strategy for targeting projections to ACC in YFP control (YFP; I) and SSFO cohorts for the retrosplenial cortex (ASP) projections (RSP->ACC; J) or AM projections (AM->ACC; K). Raw lick traces are shown for each mouse in each context on recent and remote retrieval sessions, data are mean (solid line)± s.e.m (shaded area). (L) Quantification of discrimination between high reward (HR) and aversive (A) contexts on preexpo­ sure, recent, and remote sessions for each cohort; YFP (no opsin control, N=14); ASP (RSP->ACC SSFO excitation, N=7); AM (AM->ACC SSFO excitation, N=7). *p<0.05 for YFP, ASP and AM cohorts between preexposure and remote, one-way repeated measures ANOVA with post-hoc Tukey’s multiple comparison test. Individual data points shown, with mean± s.e.m. (M) Same as in L, but for discrimination between low reward (LR) and aversive (A) contexts. *p<0.05 only for AM cohort between preexposure and remote, one-way repeated measures ANOVA with post-hoc Tukey’s multiple comparison test. Individual data points shown, with mean± s.e.m. A two-way repeated measures ANOVA was also performed and a significant interaction of Group x Context x Session was found (p = 0.0244).

To further strengthen the generality of these findings beyond the behavioral task we designed here, we also tested the effects of AM on memory consolidation in a more commonly used behavioral paradigm- contextual fear conditioning. We tested the effect of bilateral inhibition of AM-to-ACC projections during training on recent or remote recall and found a significant and isolated deficit in remote memory recall (Fig. 5E-F, 64% freezing at remote time for mCherry group vs 39% for GTACR, P<0.01 t-test). Taken together, these optogenetic and behavioral results are consistent with a critical requirement for AM activity during training to ensure remote memory time.

We next asked whether enhancing the natural excitability of the AM-to-ACC circuit during training could facilitate consolidation of a memory that otherwise would not be consolidated. To test this, we extended the current behavioral task to include two reward contexts, with non-overlapping sets of sensory cues, where one was associated with high reward (HR) and the other with a less salient, low reward (LR) outcome (Figure 5G, Methods). We optimized this task to ensure that mice would learn the two context-associations equally but consolidate only the high reward context (Figure 5I). To facilitate activity in the AM-ACC circuit in a manner that modulates the excitability of ongoing activity without exogenously driving spiking activity, we used a stabilized step function opsin (SSFO) (Yizhar et al., 2011). We injected retroAAV-Cre bilaterally in ACC, and floxed-SSFO-eYFP (or floxed-eYFP for control) bilaterally in AM thalamus, followed by implantation of optical fibers bilaterally in AM (Figures 5H and S5B). Strikingly, we found that enhancing AM-ACC excitability during training uniquely and significantly enhanced remote memory retrieval of the low reward context, effectively recalling both HR and LR at remote time (Figures 5I-5M; AM SSFO DI = 0.7368 in low reward, P=0.0111; RSP DI = 0.0164 in low reward, P=0.4833, n.s.; YFP DI = 0.0304 in low reward, P = 0.8835, n.s,) Importantly, enhancing excitability of another ACC projecting area (RSP) did not improve memory consolidation (Figure 5J,L-M). Finally, to rule out that these effects are due to a role for AM in re-training, extinction, or reconsolidation, we repeated this experiment without testing mice at a recent timepoint and again found sustained memory recall for the low reward context in SSFO-activated but not control mice (Figures S6I-L). Taken together, these results demonstrate that the AM-to-ACC projection is not only required for successful memory consolidation, but that activity enhancement at the time of learning can profoundly facilitate consolidation of otherwise unconsolidated memories.

### A fiber bundle-based method for simultaneous multi-region imaging at cellular resolution

To gain mechanistic insight into how AM thalamus facilitates memory consolidation, we aimed to perform real-time cellular resolution neural activity imaging of AM thalamus, together with hippocampus and ACC simultaneously during learning and throughout memory consolidation. Existing cellular resolution imaging approaches powerfully enable single region deep brain imaging (Dombeck et al., 2007; Hofer et al., 2009; Ghosh et al., 2011; Aharoni and Hoogland, 2019) or large-scale cortical imaging (Sofroniew et al., 2016; Weisenburger et al., 2019; Yang and Yuste, 2017), but do not provide capabilities for imaging multiple cortical and sub-cortical regions spanning different depths (i.e., thalamus, hippocampus, and cortex) simultaneously during behavior. To provide a solution, we leveraged fiber bundle-based imaging approaches (Helmchen et al., 2001), where the inherent distance created between the animal and the objective offered a scalable solution toward multi-site imaging. We also exploited recent advances in the resolution and sensitivity of multi-core fiber bundles and CMOS sensors, respectively. We thus designed a multi-region imaging system where light from multiple fiber bundles (Fujikura, 600 um Diameter, 3.3 um core to core distance) are collected onto a common objective (Nikon 10x, 0.5 NA) and projected onto a camera sensor (Photometrics Prime 95b, 95% QE), while optimizing sensitivity, resolution, and robustness to motion-related artifacts from a head-fixed or freely moving animal (Figure 6A, Methods). The resulting system was first characterized by imaging sub-diffraction size beads, where reconstruction of the PSF revealed a lateral resolution of ~5-10um, comparable to the widely used mini-scope approach (Figure S7A) (Glas et al. 2019). We further validated the ability of our system to resolve individual cells using resolution test targets (Thorlabs, Figure S7B). To image in the behaving mouse, we optimized GCaMP injection titers, GRIN-lens based surgical implantations, and the design of an adapter between the animal/GRIN lens and the fiber bundle system (Autodesk Fusion 360, Figure 6B), which together provided stable cellular resolution imaging, and access to the same neurons from the AM, HPC and ACC for up to 34 days (Figures 6C-E, Figure S7D-F). We further assessed and found that the signal-to-noise of neural activity detected from the fiber bundle system was comparable with state-of-the art wide-field recordings using matched sources from the same field-of-view (Figure S7C). Finally, although we recorded in head-fixed rodents due to the nature of our task, our adaptors provide a stable connection between the GRIN lens and the fiber bundle, and is thus compatible with studies in freely-moving animals. Thus, we developed an optical system, that interfaces seamlessly with the behaving animal, to enable high resolution imaging from multiple brain regions simultaneously and chronically during behavior (Supplementary Video 1).

**Figure 6.**
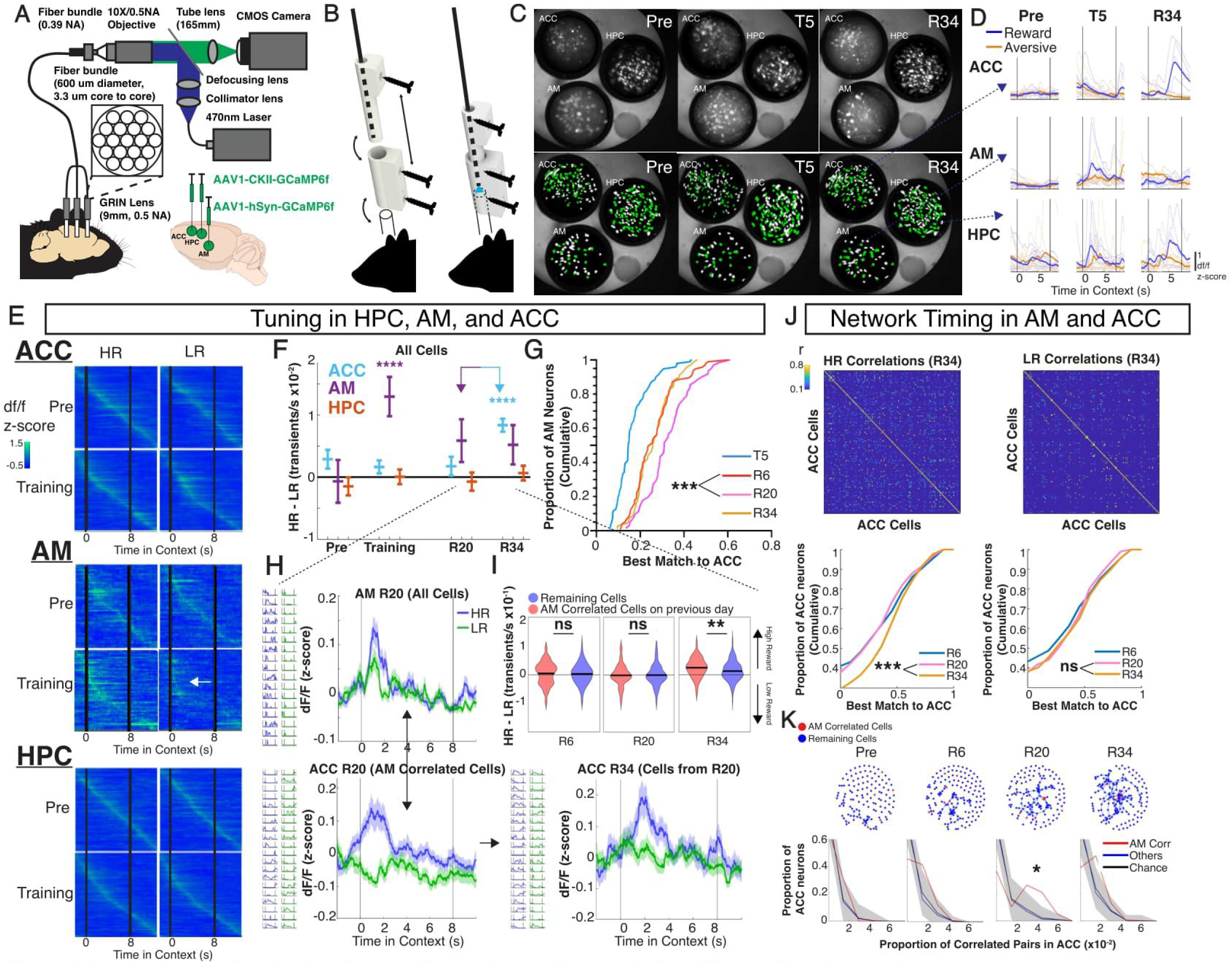
Longitudinal multi-region imaging reveals neural dynamics that distinguish consolidated memories. (A) Multi-region cellular resolution imaging setup. Fluorescence emission from the fiber bundle is collected into a common objective and projected onto a CMOS camera (see methods). (B) Custom-built apparatus for stabilizing and aligning the fiber bundle-to-GRIN lens connection. (C) Neural sources are tracked simultaneously in anterior cingulate cortex (ACC), hippocampus (HPC), and anteromedial thalamus (AM) for up to 34 days. Shown are neural sources from one mouse, same field of view tracked across preexposure (Pre), training day 5 (T5) and remote retrieval (R34). Sources captured on all days are highlighted in green. (D) Individual and mean z-scored dF, recorded for the same cells in ACC (top), AM (middle), and HPC (bottom), aligned to cue zone entry, and tracked from pre-exposure through training and remote retrieval. Dashed lines represent the source from 6C of which the dF traces are tracked. Scale: 1 df/f(z-scored) (E) Mean z-scored dF/F for every recorded cell, aligned to cue-zone entry in high reward (HR; left) and low reward (LR; right) in preexposure and training (N=3 mice). Number of neurons for preexposure and training, respectively: ACC: n=426 and n= 334; AM: n= 94 and n=75; HPC: n=461 and n=457. (F) Tuning to high reward vs low reward contex1s (see methods) for every recorded cell in ACC, HPC, and AM across days. ****p<0.0001 for training (AM), R20 (ACC) and R34 (ACC), one sample I-test. (G) Cumulative distribution of the best match correlation of ACC cells paired with AM cells across days. ***p<0.001, Wilcoxin rank-sum between R6 and R20. (H) AM and ACC correlations on R20 predict tuning in ACC on R34. Top left: cue-zone-aligned mean z-scored dF/F of all AM cells on R20 (n=111 cells). Bottom left: Mean z-scored response of ACC cells which form highly correlated pairs with at least one AM cell (n=56 cells across 3 animals, 60% correlated to one AM cell, 20% correlated to two AM cells, and 14% correlated to three or more AM cells). Bottom right: mean response of these same tracked ACC R20 cells on R34. For each mean dF/F trace, representative cells selected from one animal are shown to the left. (I) Distribution of tuning for ACC cells correlated to at least one AM cell on the previous recording session compared to the tuning of all remaining cells. •• p<0.01, Wilcoxin rank-sum test. (J) Top: Representative correlation matrices within ACC for either HR or LR context, taken from a single animal. Bottom: Cumulative distribution of best match correlation of ACC cell pairs across recording sessions for HR (left) and LR (right). ***p<0.001, Wilcoxin rank-sum between R6 and R20. (K) Top: Example from one animal of undirected network graphs of ACC cell correlations. Each node represents an individual ACC cell, and connecting edges represent significant correlations in HR (Pairwise Pearson’s r>0.3). Red nodes are the ACC cells most temporally correlated with AM cells (top 5% of Pearson’s r) shown in red. Bottom: Quantification of the proportion of pairwise correlations (number of edges for a given cell normalized by the total number of cells for a given recording secession) in ACC for either red nodes (ACC cell that is highly correlated to at least one AM cell) and blue cells (all remaining ACC cells), compared to chance (in black, see methods). Shaded area represents the 95% confidence interval around chance distribution. Significance is indicated by bins above this level.

### Anteromedial thalamus facilitates long-term contextual memory representations in cortex

We generated a cohort of mice that were injected with GCaMP6f and implanted with GRIN lenses in AM, HPC, and ACC (Figure S7G, Methods), and confirmed normal physiological activity of neurons in all regions (Figure S7H). For targeting ACC, we ensured a field of view primarily consisting of cingulate cortex, rather than the nearby M2 motor cortex, by targeting a field of view as close to the midline as possible, and confirming that motor related signals from neurons in this region are minimal (Figure S7G). For targeting AM, we ensured that ACC projecting neurons were contained within the imaging field of view, thus minimizing imaging of nearby thalamic nuclei. Implanted mice learned the previously described memory consolidation task well and showed no significant behavioral deficits compared to surgically naïve mice (Figure S7I). We recorded from all three regions simultaneously during pre-exposure (Pre), training (T1, T5), and during retrieval at recent, intermediate, and remote timepoints (R6, R20, R34). Examining all recorded cells, we observed that even as early as during training, AM cells preferentially encoded the HR compared to the LR context, a difference that was noticeably absent in HPC and ACC cells (Figure 6E and F). However, by R34, ACC, but not HPC or AM, exhibited preferential tuning to HR.

To understand how HR contextual representations progressively re-organize across these regions, we assessed inter-regional activity patterns occurring over the timescales of systems consolidation. Interestingly, we found that context-specific correlations between AM and ACC activity progressively increased across weeks (Figure 6G), in a manner that was not observed between HPC and AM or HPC and ACC (Figure S8C), and were not due to any appreciable changes in overall event rates in either AM or ACC (Figures S8A and S8B). Thus, the progressively increasing AM-to-ACC network interactions, peaking at R20, suggested potential stabilization of salient contextual representations from AM to ACC between R20 and R34. To test this, we identified ACC cells that were highly correlated with AM cells on R20 (Fig. 6H top and bottom left) and determined their tuning on R34. We found notably that such ACC cells identified on R20 were significantly more likely to maintain their tuning to the HR context on R34, compared with remaining ACC cells (Figures 6H-I and S8D, Wilcoxin rank-sum P<0.01), or with ACC cells tuned to HPC on R20 (Figure S8E-F, Wilcoxin rank-sum p=0.6175).

As a signature of consolidation, in addition to increasing tuning to HR, we observed progressively increasing intra-ACC ensemble correlations over weeks (R6 to R34) specifically in the HR but not LR context (Figure 6J). Notably, cells in ACC that had high functional correlations with AM were significantly more likely to be “central” in the network than one would expect by chance (Figure 6K) and again specifically on R20. This was not the case for cells in ACC having high functional correlations with HPC (Figure S8G). These data suggested that AM may facilitate not only the tuning of ACC contextual representations, but also their temporal synchrony within ACC, at remote time.

Taken together, these results suggest a model in which AM performs early selection of salient memories during training, and facilitates their re-organization to cortex via progressive increases in <u>inter</u>-regional functional coupling between AM and ACC (that peaks at R20), thus coordinating the stability and synchrony of memory ensembles within ACC at remote time (that peaks at R34). The selection of salient memories by AM in training provides an explanation for how optogenetic manipulations (either inhibition or excitation) during training may still impact circuits at remote time points. We tested this model explicitly by developing another cohort of mice with bilateral stGtACR2-based inhibition of AM-to-ACC projections during training (T1-T5) while imaging in ACC at recent and remote time (Figures 7A and S9A). We found indeed that inhibition of AM-to-ACC activity during training again disrupted memory recall at remote but not recent time points (Figure S9C), and that this was associated with a significant but modest loss of contextual tuning (Figure 7B), but more strikingly, a severe disruption in context-specific ensemble timing within ACC at remote time (Figure 7C). Thus, we present a model where AM-to-ACC interactions during training are required for establishing the eventual tuning and synchronicity of cortical contextual representations associated with successful consolidation and remote memory recall (Figure 7D).

**Figure 7.**
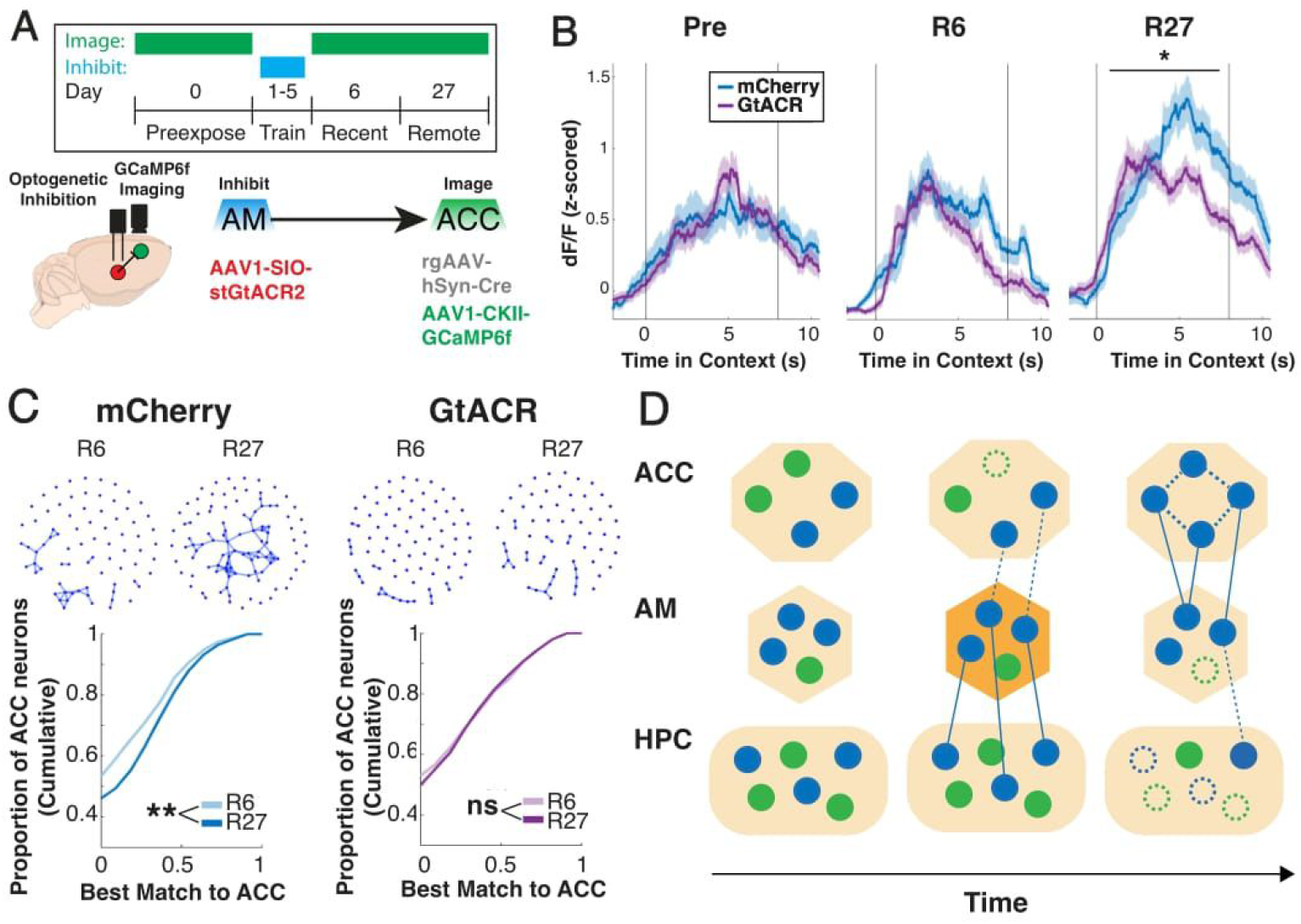
AM is required to stabilize the tuning and timing of long-term contextual representations in cortex. (A) Experimental design for optogenetic inhibition and imaging experiment. rgAAV1-hSyn-Cre and AAV1-CKII-GCaMP6f was injected into anterior cingulate cortex (ACC) and AAV1-SIO-st­ GtACR2 was injected into anteromedial thalamus (AM). Imaging was performed on preexposure, recent and remote retrieval sessions, whereas inhibition occurred during each training session. (B) Mean z-scored dF for context-tuned neurons, aligned to cue-zone entry, compared between mCherry and GtACR mice on preexposure, recent (R6) and remote (R27) retrieval. *p<0.05, two-sample I-test. (C) Top: Examples of undirected network graphs of ACC cell correlations in HR from mCherry (left) and GtACR (right) mice during recent and remote retrieval. Each node corresponds to a cell in ACC and each edge is a highly correlated cell pair (Pairwise Pearson’s r>0.3) Bottom: Cumulative distribution of the best-match correlation for every cell in HR from recent to remote retrieval. N=5 mCherry vs 7 GtACR mice. **P<0.01, Wilcoxin Rank-Sum test. (D) Proposed model for how AM selects and stabilizes consolidated memory representations in cortex. Selection of information about salient cues by AM leads to coordinated responses in ACC at remote timepoints, eventually allowing for recall that is independent of HPC.

## DISCUSSION

In this study, we demonstrate that the AM thalamus serves as a critical gateway for the consolidation of memories to long-term cortical storage. Four major observations support this conclusion: 1. Bulk activity recordings revealed neural correlates of a learned association in anterior thalamus, but not amygdala or entorhinal cortex, that persisted for several weeks during the consolidation window; 2. Bilateral inhibition of AM thalamus to anterior cingulate projections during training disrupted memory consolidation and remote recall; 3. Gain amplification of the AM thalamus to anterior cingulate projections, but not another retrosplenial-to-cingulate projection, during training was sufficient to enhance consolidation of otherwise unconsolidated memories, and 4. Multi-region cellular resolution imaging identified that while hippocampus encodes multiple memories equally, AM preferentially encodes salient experiences during training and establishes inter-regional correlations with cortex during consolidation, which are required for coordinated and stable context representations in ACC at remote time for behavioral consolidation.

Overall, we find that while both hippocampus and cortex form early representations of context, there is increasing dependence on cortex over time (Do-Monte et al., 2015; Kitamura, T. et al., 2017; DeNardo et al., 2019) that is shaped critically by thalamo-cortical interactions. Furthermore, with increasing dependence on cortex, we find a concomitant decrease in hippocampal dependence, consistent with an active reorganization of memory representations across the brain toward progressively more stable forms. However, it is likely that for memories that are highly spatial, detailed, or autobiographical, the hippocampus continues to remain engaged (Nadel and Moscovitch, 1997; Yonelinas et al., 2019; Barry and Maguire, 2019). Interestingly, the preferential encoding of salient experiences by AM has been described more generally in thalamus (Zhu et al., 2018) and is further supported by evidence of known inputs to AM from reward-processing regions (Harris et al., 2019). Thus, AM thalamus’ functions in encoding of salient experiences, together with its role in temporally coordinating ACC ensembles, offers a suitable mechanism for the consolidation of select memories to long-term storage, where aspects of this model have also been computationally predicted (Tomé et al., 2020; Fiebig and Lansner, 2014). Notably, patients with lesions to the anterior thalamus can present with varying degrees of intermediate to retrograde amnesia (Aggleton et al. 1999, Miller at al., 2001). In particular, Korsakoff Syndrome patients with bilateral lesions of the medial mammillary to anterior thalamus to cortex pathway present with severe, and sometimes isolated, retrograde amnesias.

We note that the very nature of longitudinal imaging used in this study, while useful for tracking the changing nature of memory representations across the brain over time, requires repeated behavioral measures of recall. We cannot therefore exclude the possibility that the changes in neural activity patterns that we observe are partially due to extinction or reconsolidation. Thus, in the future, a systematic use of separate cohorts for each terminal recall time point would fully address any confounds due to reconsolidation. Furthermore, while we focused here on the contributions of anteromedial thalamus, there may be interesting crosstalk with other thalamic nuclei (Do-Monte et al., 2015; Vetere et al., 2021), and limbic circuits (Kitamura et al., 2017), which collectively may provide coordinated or separate routes of consolidation for different types of memories. It will also be important to determine how neural plasticity in these circuits operate together with neural oscillations (Wilson and McNaughton, 1994; Montgomery et al., 2008; Karlsson and Frank, 2009) during online and offline periods (Latchoumane et al., 2017; Goto et al., 2021; Chouhan et al., 2020) to enable stable long-term representations. This work also provides new circuit level entry points toward the identification of transcriptional and epigenetic programs (Crick, 1984; Rajasethupathy et al., 2012; Gräff et al., 2014; Chen et al., 2020; O’Banion, et al. 2020) enabling the maintenance of long-term memories. These and related approaches will enable a deeper mechanistic insight into the mysteries of slow brain re-organization underlying memory consolidation.'

## ACKNOWLEDGMENTS

We thank the Rockefeller Precision Instruments & Fabrication Facility for help with design and implementation of the fiber adaptors. We thank Andrea Terceros for sharing unpublished data related to the development and extensions of the behavioral task described here. We thank Chelsea Noble for assistance with stereotactic surgeries. We are grateful to Charles Gilbert, Cori Bargmann, Leslie Vosshall and Rajasethupathy lab members for helpful discussions related to various aspects of the study. This work was supported by a Medical Scientist Training Program grant from NIGMS of the NIH (T32GM007739) to the Weill Cornell/Rockefeller/Sloan Kettering Tri-Institutional MD-PhD Program, and an NRSA fellowship awarded to AT, an HHMI Gilliam fellowship awarded to JMR, and grants from the Mathers foundation, Klingenstein foundation, and from the National Institutes of Health under award number DP2AG058487 to PR.

## AUTHOR CONTRIBUTIONS

ACT, JMR, and PR conceived the study and designed the experiments. ACT characterized the fiber bundle imaging system, performed all imaging experiments, developed associated post-processing pipelines and analyzed the data. ACT also performed paired optogenetic & imaging experiments. JMR developed and characterized the VR-based behavioral task, and performed photometry and optogenetics experiments and associated analyses. YR assisted with the development of GRIN lens stereotactic surgeries and VR-based behaviors. AT performed the fear-conditioning experiment. NY assisted in population level analyses for imaging data. SK developed the first version of the fiber-bundle imaging system. SS assisted with VR based behaviors and histology. FH performed anatomical tracing studies. ABO assisted with project discussions and coordination. ACT and PR wrote the manuscript with input from all authors. PR supervised all aspects of the work.

## DECLRATION OF INTERESTS

The authors declare no competing financial interests.

## STAR METHODS

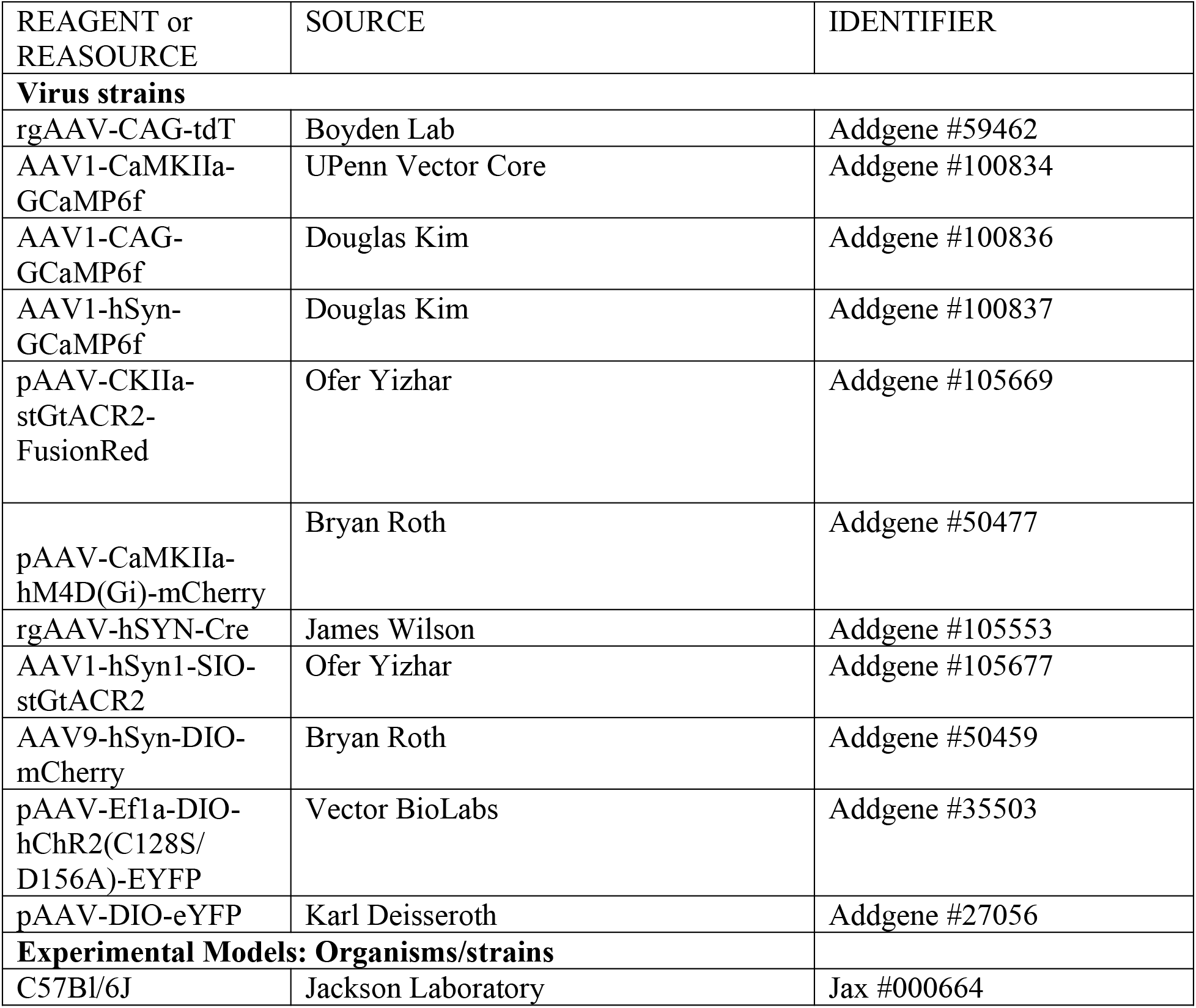

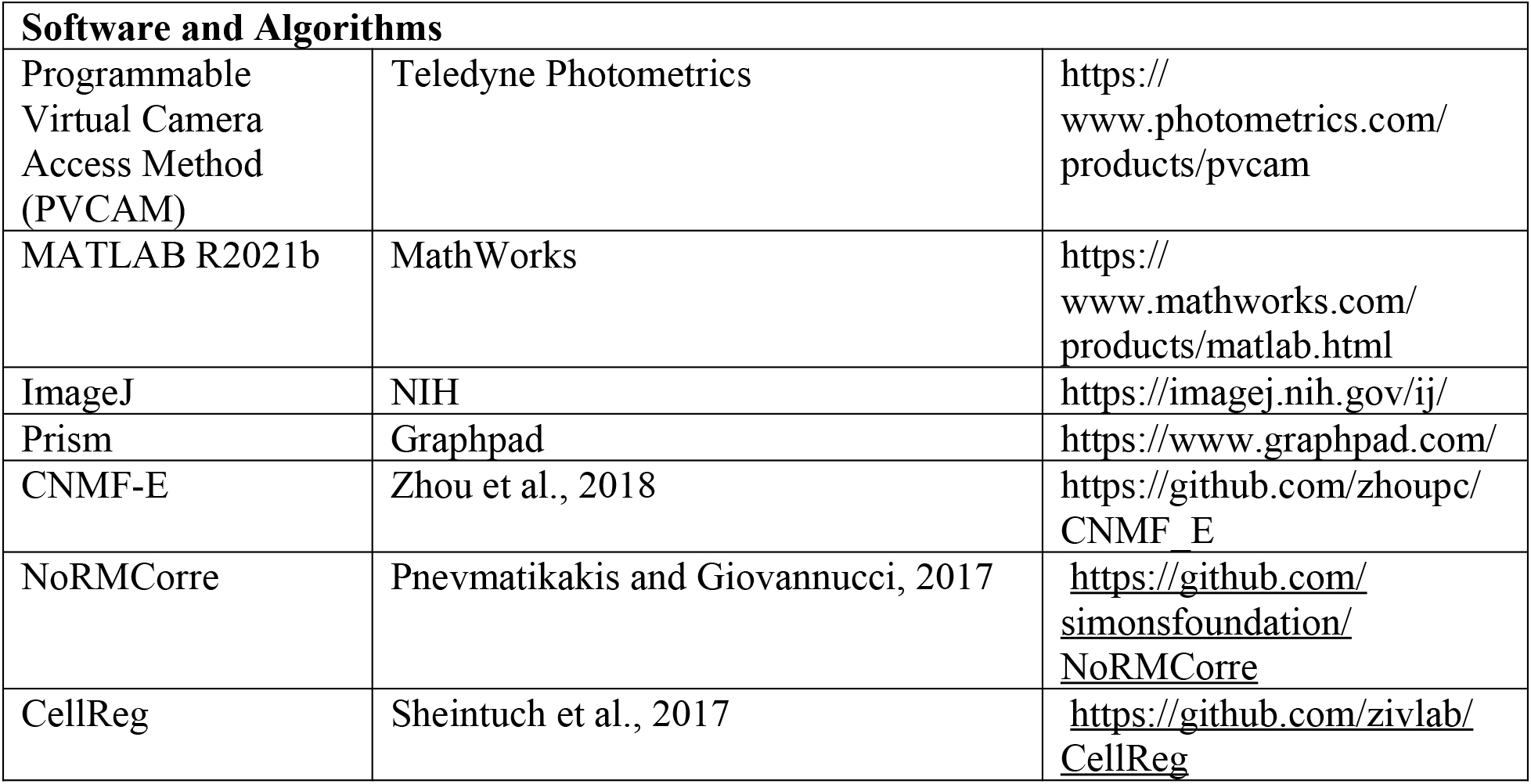

## EXPERIMENTAL MODEL AND SUBJECT DETAILS

### Mice

All procedures were done in accordance with guidelines derived from and approved by the Institutional Animal Care and Use Committees (protocol #19112H) at The Rockefeller University. Animals used were 8-10 weeks-old wild-type male and female (counterbalanced across experimental conditions when applicable) C57BL/6J mice (Jackson Laboratory, Strain #000664) at the time of surgery. Mice were group housed (3-5 per cage) with ad libitum food and water unless mice were water restricted for behavioral assays, in which case they were given 1 mL water a day. Body weight was monitored daily to ensure it was maintained above 80% of the pre-restriction measurement. Surgical procedures and viral injections were carried out in mice under protocols approved by Rockefeller University IACUC and were performed in mice anesthetized with 2% isoflurane using a stereotactic apparatus (Kopf).

## METHOD DETAILS

### Surgical Procedures

Puralube vet ointment was applied to the eyes and 0.2mg/kg meloxicam was administered intraperitoneally using a 1mL syringe. Hair from the scalp was trimmed, and the area was sterilized using povidone-iodine swabs and subsequently ethanol swabs. An incision covering the anteroposterior extent was made to allow access to the skull. Injection sites were accessed using a dental drill which made 0.5mm holes through the skull. All virus was injected using a 35G beveled needle in a 10ul NanoFil Sub-Microliter Injection syringe (World Precision Instruments) controlled by an injection pump (Harvard Apparatus) at a rate of 100nl/min. After all viral delivery, an additional 5-10 mins delay was applied to avoid backflush before slowly removing the injection needle. Animals that required cannulas or GRIN lenses were implanted immediately following viral injection. Following surgery, mice were allowed to recover in a single housed cage for up to 12 hours, and were given meloxicam tablets. Mice were typically housed for three weeks to allow for adequate expression before behavioral testing or histology.

#### Viral injections

- In retrograde tracing experiments, mice were unilaterally injected in ACC (A/P +1.0, M/L, ±0.35, D/V −1.4) with rgAAV-CAG-tdT at a volume of 500 nl (1.0 × 10^13^ vg/mL).
- For multi-fiber photometry experiments, 1ul of AAV1-CaMKIIa-GCaMP6f (UPenn Viral Core, diluted to 5 × 10^12^ vg/mL) was injected into ACC, HPC (A/P: −1.5, M/L: ±1.5, D/V: −1.6), ENT (A/P: −4.35, M/L: ±3.75, D/V: −4.0), and BLA (A/P: −1.23, M/L: 2.75, D/V: −4.7) while AAV1-CAG-GCaMP6f was injected into ANT. One week after virus injection, mice were unilaterally implanted with 1.25 mm ferrule-coupled optical fibers (0.48 NA, 400 μm diameter, Doric Lenses) cut to the desired length so that the implantation site is ~0.2 mm dorsal to the injection site.
- For cellular imaging, AAV1-CaMKIIa-GCaMP6f (5.0 × 10^12^ vg/mL) was injected into ACC and HPC while AAV1-hSyn-GCaMP6f was injected into AM (5.0 × 10^12^ vg/mL)
- For optogenetic inhibition of HPC, AAV1-CaMKIIa-stGtACR2 (1 × 10^13^ vg/mL) was injected into HPC bilaterally. For controls, AAV1-CaMKIIa-mCherry (7 × 10^12^ vg/mL) was injected.
- For chemogenetic inhibition of HPC and ACC, AAV9-CaMKIIa-hM4D(Gi) (1 × 10^13^ vg/mL) was injected into ACC or HPC bilaterally. For controls, AAV9-CaMKIIa-mCherry (1 × 10^13^ vg/mL) was injected bilaterally in either region.
- For optogenetic inhibition of AM-ACC projections, rgAAV-hSYN-Cre (1.20 × 10^13^ vg/mL) was injected into ACC bilaterally and either AAV1-hSyn1-SIO-stGtACR2 (1.50 × 10^13^ vg/mL) or AAV9-hSyn-DIO-mCherry (9.0 × 10^12^ vg/mL) for controls was injected bilaterally into AM.
- In SSFO experiments, rgAAV-hSyn-Cre (1.3 × 10^13^ vg/mL) was injected bilaterally in ACC and AAV-Ef1a-DIOhChR2(C128S/D156A)-EYFP (Vector BioLabs, 1.0 × 10^13^ vg/mL) bilaterally in AM or RSP (A/P: ±0.40, M/L:-2.0, D/V: −0.7) for behavioral control and AAV-DIO-eYFP (1.0 × 10^13^ vg/mL) into AM for experimental control.

#### TRIO tracing

For the “tracing of input-output” (TRIO) tracing of AM-ACC, CAV-Cre (500 nL, 2.5 × 10^12^ vg/mL) was injected into ACC, and a combination of AAV8-CAG-FLEX-TCB/ AAV8-CAG-FLEX-oG-WPRE-SV40 (500 nL each, 2.5×10^12^) was injected into AM. After two weeks, Env-RV-dG-eGFP (500 nL, 1 × 10^9^ vg/mL) was injected into AM. Five days later, mice were perfused and tracing was confirmed via histology.

#### Cannula implants

One week after viral injections, mice undergoing photometry or optogenetic experiments were implanted with fiber optic cannulas (Doric Lenses). For photometry, mice were unilaterally implanted with 1.25 mm ferrule-coupled optical fibers (0.48 NA, 400 μm diameter, Doric Lenses) cut to the desired length so that the implantation site is ~0.2 mm dorsal to the injection site. For optogenetics, mice were implanted bilaterally with 1.25mm cannulas (0.22 NA, 200um diameter, Doric Lenses). In both cases, cannula implants were slowly lowered using a stereotaxic cannula holder (Doric) at a rate of 1 mm/min until it reached the implantation site, 0.2 mm dorsal to the injection site. In the case of bilateral AM optogenetic inhibition, one cannula was implanted at a 10-degree angle laterally to the skull in order to prevent stereotactic hindrance.

Optic glue (Edmund Optics) was then used to seal the skull/cannula interface and a custom titanium headplate was glued to the skull using adhesive cement (Metabond).

#### GRIN lens implants

Immediately following viral injections, mice undergoing calcium imaging were implanted with gradient-index (GRIN) lens(es). An incision covering the anteroposterior extent was made, and the skin overlying the skull was cleared. The skull was then cleared and textured using a scalpel. Using a dental drill, 1mm diameter holes were made at stereotactically determined sites of implantation. Site of drilling was immediately covered using chilled 1x PBS, and using a sterile 28 G x 1.2” insulin syringe and low-pressure vacuum suction, the underlying dura was removed. GRIN lenses (0.6 mm diameter, 7.2mm length, 0.5 NA from Inscopix) were wrapped in a 0.68mm wide custom length stainless steel sleeve (McMaster, catalog # 8987K54) using optic glue, made to cover only the part of the lens held external to the brain. With a 0.5mm burr (Fine Science Tools) attached to a stereotaxic cannula holder, the GRIN was slowly lowered into the brain at a rate of 1mm/min, ending 0.2mm dorsal to the injection site. The skull was constantly flushed with chilled 1x PBS. Every time the lens moved 0.8 mm more ventral, it was temporarily retracted 0.4 mm dorsally at the same rate, before continuing down again. We found this especially helpful to maximize the number of observed cells when imaging in very deep regions. The skull-sleeve connection was then sealed with glue, and further secured with adhesive cement. For multi region imaging, the two other GRIN lenses were implanted in the same manner, with consideration to stereotactic hindrance from each additional lens (First AM, then ACC, and finally HPC). A custom titanium headplate was glued to the skull using adhesive cement. Immediately following surgery, mice were injected with 0.2mg/kg dexamethasone subcutaneously to reduce inflammation.

### Histology

Animals were deeply anesthetized with 5% isoflurane before transcardial perfusion with ice-cold PBS and 4% paraformaldehyde in 0.1M PB. Brains were then post-fixed by immersion for ~24 hours in the perfusate solution followed by 30% sucrose in 0.1M PB at 4°C. The fixed tissue was cut into 40 μm coronal sections using a freezing microtome (Leica SM2010R), free-floating sections were stained with DAPI (1:1000 in PBST), and mounted on slides with ProLong Diamond Antifade Mountant (Invitrogen). Images were taken on a Nikon Inverted Microscope Eclipse Ti-E with a 4x/0.2 NA objective lens. Whole-slide-images were stitched with NIS-Elements imaging software and further analyzed in ImageJ and MATLAB.

### Virtual Reality System

We used a custom-built virtual reality environment, modified from a previously reported version (Rajasethupathy et al., 2015). In brief, a 200-mm-diameter styrofoam ball was axially fixed with a 6-mm-diameter assembly rod (Thorlabs) passing through the center of the ball and resting on 90° post holders (Thorlabs) at each end, allowing free forward and backward rotation of the ball. Mice were head-fixed in place above the center of the ball using a headplate mount. Virtual environments were designed in the virtual reality MATLAB engine ViRMEn (Aronov and Tank, 2014). The virtual environment was displayed by back-projection onto white fabric stretched over a clear acrylic hemisphere with a 14-inch diameter placed ~20 cm in front of the center of the mouse. The screen encompasses ~220° of the mouse’s field of view and the virtual environment was back-projected onto this screen using a Vamvo Ultra Mini Portable projector. The rotation of the styrofoam ball was recorded by an optical computer mouse (Logitech) that interfaced with ViRMEn to transport the mouse through the virtual reality environment. A National Instruments Data Acquisition (NIDAQ) device was used to send out TTL pulses to trigger the CMOS camera, laser for optogenetics, and the various Arduinos controlling tones, odors, airpuff, lick ports. Additionally, the NIDAQ recorded the capacitance changes of the lick port when licking occurred and the CMOS camera exposures to align lick rate and neural recording/imaging to trial events.

### Behavioral shaping

Starting approximately 3 weeks after mice recovered from surgery, they were put on a restricted water schedule, receiving 1 mL of water in total per day. Body weight was monitored daily to ensure it was maintained above 80% of the pre-restriction measurement.

After a week of water deprivation, mice were habituated to the styrofoam ball for 2 days by receiving their 1 mL of water per day head-fixed. Then mice were put onto a linear track (vertical gray bars) where water release was contingent on walking a short distance to the outcome zone (white horizontal bars) where they received 5 seconds of water delivery. Over the course of a session, and in subsequent days, the distance needed to travel to enter the outcome zone increased. If a mouse took longer than 10 minutes to receive their 1 mL of water on a given day, the distance needed to travel to get water was repeated on the following day until they could reliably walk on the ball for water under 10 minutes. Once mice could walk down the full length of the linear track used in training, we introduced the inter-trial interval and a timer (5 seconds for photometry experiments, 8 seconds for optogenetics and single-cell imaging experiments) to encourage mice to actively engage with the task. Once all mice from a cohort were able to complete at least 80% of initiated trials, training began.

### Behavioral task

In the final version of the task that was used during photometry, optogenetics, and imaging experiments, mice ran down a virtual linear track to reach an outcome zone and used contextual cues presented along the track to predict the reinforcement they will receive (~4ul of sucrose water or airpuff). At the beginning of the linear track, mice self-initiate sessions by running through a start zone above a set speed of 12 cm/s, (~20% of the ball perimeter in a second). Visual, auditory, and olfactory cues were then presented in succession 1 second after one another. The visual cues consisted of blue stars or yellow triangles designed within the ViRMEN GUI (Aronov and Tank, 2014) and presented along the walls of the linear track. The auditory cues consisted of 5 KHz or 9 KHz tones outputted by a thin plastic speaker (Adafruit) and olfactory cues consisting of ɑ-pinene or octanol were diluted with mineral oil to 10% and released by a custom-built olfactometer. Both auditory and olfactory cues were outputted by Arduino code under the control of ViRMEN code. The cues for reward context were blue stars, 5KHz tone, and alpha-pinene while the cues for aversive context were yellow triangles, 9KHz tone, and octanol. For photometry experiments, all cues were turned off after 5 seconds into the context zone. For optogenetics and single-cell imaging experiments, the cue zone was extended to last 8 seconds. Entry into the outcome zone was contingent upon running down a linear distance of 60 cm along the ball within the cue zone time limit. If that distance was not reached, mice would be placed in a 5 second inter-trial interval proceeding the next trial. The outcome zone consisted of free access to 10% sucrose water presented by a lickometer (reward context) or a 50ms airpuff presented by a pipette tip cut to a fine point 3 inches away from the snout (aversive context). Both reinforcements were outputted by Arduino code under the control of ViRMEN code. After the outcome zone mice were transported to a 5 second inter-trial interval before starting the next trial.

Performance in the task was assessed by average speed (during the first 5 seconds in cue zone) and average anticipatory lick rate (during the last 2 seconds in the cue zone) for all reward and aversive context trials in a given session. Prior to training, mice were given a “pre-exposure” session where they were exposed to each set of cues, with tap water given upon reaching the outcome zone in both. They were then given 5 days of training (referred to as T1-T5), where reinforcement was given in the outcome zone. For the photometry experiment, mice began 5 minutes of retrieval sessions where no reinforcement is given on days 6, 13, 20, 27, 41, and 55 since training began. After each retrieval session, mice underwent 8 trials of each context in a “retraining” session where reinforcement was given again, to avoid memory extinction and loss of engagement in the task. For the optogenetics and imaging experiments with early and remote retrieval sessions, mice were tested with sessions consisting of 4 interspersed trials of each context type (for a total of 12 trials), followed by a re-training session of the same length. For experiments with only remote retrieval, mice received a single session again consisting of 4 trials of each context.

### Modified behavior with three contexts

Mice undergo behavioral shaping as outlined above. During pre-exposure and training days, a third low reward (LR) context with non-overlapping cues (green diamond visual cues, 11 kHz tone, and orange-scented essential oil diluted in mineral oil) was included alongside high reward (HR) (same as “R” context in two-context task) and aversive (A) contexts. Presentation was randomized. In low reward training trials, 10% sucrose water was delivered through the lickport at a rate of 1 drop every second for five seconds. No animal was given more than 1mL of sucrose on a given day, and each session concluded after the completion of 8 trials per context. In SSFO excitation experiments, mice were trained on the task as outlined above for 5 days, then moved onto retrieval sessions on days 6 and 27. For multiple-region GRIN imaging experiments, mice had retrieval sessions on days 6, 20, and 34. Like in other trials, mice exposed given brief re-training sessions with reinforcement to minimize extinction.

### Behavioral analysis

In all behavioral experiments, we assessed learning and memory recall by calculating a normalized lick rate difference, which we refer to as the discrimination index (DI). The DI was calculated as follows:

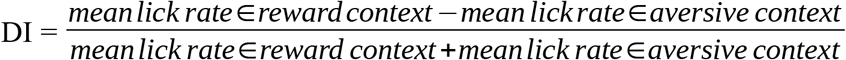

A DI of 1 therefore indicates perfect discrimination, while a DI of 0 indicates chance performance. For training sessions, the lick rate was only calculated in the window of time 4 seconds preceding the onset of the outcome zone. On retrieval sessions, we included lick rates during the outcome zone, as no reinforcement is provided but mice still expressed anticipatory licking behavior. Repeated measures ANOVA with Tukey’s post hoc test was used to assess learning on later days by comparing to discrimination during pre-exposure.

### Optogenetic inhibition of hippocampus

Mice were injected with AAV1-CaMKII-stGtACR2 bilaterally in HPC, while control cohorts were injected with AAV1-CaMKII-mCherry. Cannulas were implanted directly above the injection sites. After three weeks, mice underwent shaping as described above, then moved onto training. For inhibition during training, light from a 473nm laser (15 mW at fiber tip) was delivered through a mono fiber optic patch cord for 13 seconds (cue zone followed by reinforcement zone) upon the animal entering the cue zone, throughout the duration of training (~15 minutes). For mice with optogenetic inhibition during retrieval, we administered optogenetic stimulation throughout all retrieval sessions from the beginning of cue zone entry to the end of outcome zone entry.

### Chemogenetic inhibition of hippocampus and ACC

For chemogenetic silencing experiments, we injected AAV9-CaMKIIa-hM4D(Gi) (or AAV9-CaMKII-mCherry for controls) bilaterally into either hippocampus or ACC. For a week prior to the remote retrieval session, mice were habituated to handling and intraperitoneal injections of saline. A solution of clozapine N-oxide (CNO) was prepared at a concentration of 0.5 mg/mL, and mice were injected at a dosage of 5 mg/kg. Behavioral experiments were conducted 45 minutes after injection.

### In Vivo Multi Site Photometry Recordings

#### Photometry Setup

A custom multi-fiber photometry setup was built as previously (Kim et al. 2016). with some modifications that were incorporated to increase signal to noise, detailed below. Excitation of the 470 nm (imaging) and 405 nm (isosbestic control) wavelengths were provided by LEDs (Thorlabs M470F3, M405FP1) which are collimated into a dichroic mirror holder with a 425 nm long pass filter (Thorlabs DMLP425R). This is coupled to another dichroic mirror holder with a 495 nm long pass dichroic (Semrock FF495-Di02-25×36) which redirects the excitation light on to a custom branching fiberoptic patchcord of five bundled 400 mm diameter 0.22NA fibers (BFP(5)_400/430/1100-0.48_3m_SMA-5xMF1.25, Doric Lenses) using a 10x/0.5NA Objective lens (Nikon CFI SFluor 10X, Product No. MRF00100). GCaMP6f fluorescence from neurons below the fiber tip in the brain was transmitted via this same cable back to the mini-cube, where it was passed through a GFP emission filter (Semrock FF01-520/35-25), amplified, and focused onto a high sensitivity sCMOS camera (Prime 95b, Photometrics). The multiple branch ends of a branching fiberoptic patchcord were used to collect emission fluorescence from 1.25mm diameter light weight ferrules (MFC_400/430-0.48_ZF1.25, Doric Lenses) using a mating sleeve (Doric SLEEVE_ZR_1.25). The excitation is alternated between 405nm and 470nm by a custom made JK flip flop which takes the trigger input from the sCMOS and triggers the two excitation LEDs alternatively. Bulk activity signals were collected using Photometrics data acquisition software, Programmable Virtual Camera Access Method (PVCAM).

#### Photometry Recordings

While mice performed the VR-based contextual discrimination task we recorded bulk calcium signals from five regions: ACC, HPC, ANT, BLA, and ENT simultaneously. We recorded at 18 Hz with excitation wavelengths alternating between 470 nm and 405nm, capturing calcium dependent and independent signals respectively, resulting in an effective frame rate of 9 Hz.

#### Data Processing

For analysis, the images captured by the CMOS camera were post-processed using custom MATLAB scripts. Regions of interest were manually drawn for each fiber to extract fluorescence values throughout the experiment. The 405-nm reference trace was scaled to best fit the 470-nm signal using least-squares regression. The normalized change in fluorescence (dF/F) was calculated by subtracting the scaled 405-nm reference trace from the 470-nm signal and dividing that value by the scaled 405-nm reference trace. The true baseline of each dF/F trace was determined and corrected by using the Matlab function *msbackadj*, estimating the baseline over a 200 frame sliding window, regressing varying baseline values to the window’s data points using a spline approximation, then adjusting the baseline in the peak range of the dF/F signal.

In order to ensure signal fidelity, we excluded individual regional recordings with poor signal-to-noise ratios. A Power Spectral Density (PSD) estimate was calculated for each recording across the entire session. Because low fidelity signals are associated with high frequency noise components (Figure S3B), a noise estimate was calculated as the PSD in the range of 0.25-0.5x the sampling rate. Peak-to-noise ratio (PNR) was calculated by dividing the max dF/F of this PSD estimate. Plotting the distribution of all signal PNR revealed an empirically defined cluster of low PNR recordings below a value of 7 (Figure S3C), which accounted for 6.55% of all recordings, spread out across regions and days. These were subsequently excluded from further analysis.

#### Bulk neural responses

The adjusted calcium signals from Photometry were aligned to task events (for example, cue presentation, airpuff, lick rate, etc) in ViRMEn by time-stamping behavioral frames captured through the NIDAQ. Photometry signals from all animals from a given region were Z-scored across the entire session. The mean regional responses to task variables (Figures 3C, 4A, and 4C), is the mean of these aligned Z-scored signals across all animals, with s.e.m. calculated across all recorded trials. We then sought to quantify the difference in temporal divergence activity patterns observed in either context. Such differences have been previously described to be similar in magnitude whether mice learn a rewarding and aversive context as to when they learn either in comparison to an otherwise neutral context (Yadav et al., 2022). To calculate the differential response to context entry (Figure 4B), for each reward or aversive trial, we calculated the change in Z-scored signal from 0 to 2 seconds into the cue zone. For example, this differential signal may be significant if, on average, the photometry signal in one context moves in either direction (more or less positive) relative to the other context upon entry into the cue zone. To calculate the differential anticipatory response (Figure 4D), the summed mean difference between the signal across all reward and fear trials in the anticipatory window (2.5 seconds prior to the end of the cue zone) was compared to the summed mean difference between the signal across both trial types in the first half of the cue zone (0 - 2.5 seconds into the control). This measurement would be significant if the relative difference in magnitude of the context signals diverged as the mouse progressed from context entry toward the outcome zone.

#### Linear encoding model

We used a generalized linear model to estimate the encoding of various task parameters by bulk neural activity in each recorded region. In our model, Z-scored dF/F is predicted by a combination of external stimuli, trial-long events, and continuous behavior. External stimuli include the trial-start zone, context-specific cues (reward/fear visual, auditory, and olfactory cues), and training feedback (sucrose or air puff). The predictors for these stimuli were generated by convolving the time course of each with a set of 6 gaussian kernels of various sizes (2-σ width ranging from 1 to 6 seconds, see figure below)

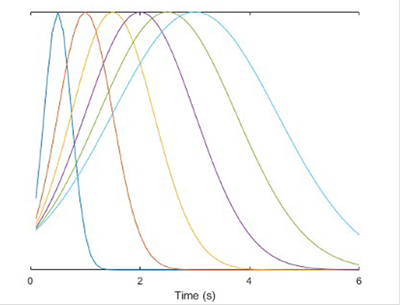

to allow for variability in the neural response. This range was determined by assessing the distribution of calcium event widths across all mice and regions (Figures S4C and S4D). The trial-long predictor, context identity of the current trial, were either 1 or 0 for all time points in the trial. Continuous behavioral predictors (speed, lick rate, and trial time) were input directly into the model. Finally, we included an L2 penalty (ridge regression) to prevent overfitting.

The full equation for the model was:

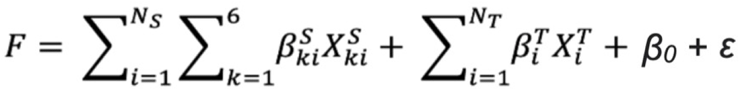

where F is dF/F for a given region, 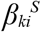 and 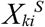 are the encoding weights and predictors, respectively, for gaussian kernel *k* and external stimulus *i*; 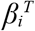 and 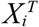 are the weights and predictors for the ith trial-long variables, and ε is the error. We fit the model on each session using 1000 rounds of 3-fold cross-validation, where the model was trained on a random split of 2/3 of trials. Accuracy was assessed by calculating the R2 between the predicted and true dF/F on the held-out trials, after concatenating across each fold.

The proportion of variance explained for a given variable was calculated adapted from previously described methods (Engelhard et al., 2019). Briefly, we first calculated the explained variance of the full model, *R^2^ _f_*. Then, a partial model was built with a given variable, *i*, excluded as a predictor and the variance explained for this partial model was calculated 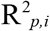. The contribution of the selected variable to the variables explained is then calculated as: 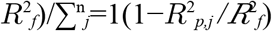.

### Optogenetic inhibition of AM Projections to ACC

Mice were injected with rgAAV-hSyn-Cre bilaterally in ACC and AAV1-hSyn1-SIO-stGtACR2 bilaterally in AM for projection specific expression of GtACR2 in AM. Control cohorts were injected with rgAAV-hSyn-Cre in ACC and AAV9-hSyn-mCherry in AM. After three weeks We continued to administer optogenetics throughout the subsequent retraining sessions, except in cohorts tested without intermediate retrieval sessions.

### Optogenetic activation of AM projections to ACC

Mice were injected with rgAAV-hSyn-cre bilaterally in ACC and pAAV-Ef1a-DIOhChR2(C128S/D156A)-EYFP bilaterally in AM for projection specific expression of SSFO in the AM. For RSP control mice, pAAV-Ef1a-DIOhChR2(C128S/D156A)-EYFP was injected into RSP and rgAAV-hSyn-cre in ACC. For optical control mice, rgAAV-hSyn-Cre was injected into ACC and pAAV-CAG-DIO-eYFP was injected into AM. Dual fiber optic cannulas were implanted in AM or RSP for the respective cohorts. At the beginning of each training session, a blue 470nm light was on for 5 seconds (at a power of 15 mW measured at the fiber tip). At the end of 30 minutes, or at the end of 10 completed trials for each context (whichever came first), a secondary pulse of yellow light at 589 nm was administered to deactivate the SSFO and return the membrane potential of transfected neurons to resting values.

### Contextual fear conditioning

Mice were injected with rgAAV-hSyn-Cre bilaterally in ACC and AAV1-hSyn1-SIO-stGtACR2 bilaterally in AM for projection specific expression of GtACR2 in AM. Control cohorts were injected with rgAAV-hSyn-Cre in ACC and AAV9-hSyn-mCherry in AM. Optical cannulas were implanted directly above the injection sites. On training day, mice were placed in Context A for a 6 minutes while 470nm light was delivered throughout. In that time they received 4 interspersed foot shocks. Mice were subsequently tested without light the following day for recent retrieval and after 3 weeks for remote retrieval. Freezing behavior was quantified as the percentage of time immobile in the first minute in Context A as recorded by a near-infrared camera.

### In Vivo Multi-Region Calcium Imaging

After surgical implants of GRIN lenses in ACC, HPC and AM, mice were given one month to recover in their home cage. Mice were selected for behavior and imaging only if multiple clear transients were identifiable in all three regions 3 weeks post surgery.

#### Microscope

A bundle of three imaging fibers with individual diameters of 0.6mm and core-to-core distance of 3.3 μm (Myriad Fibers, FIGH-30-650s) was coupled to the imaging objective (Nikon, CFI Fluor 10X, NA=0.5). The objective path was then passed through a dichroic filter (Semrock FF01-520/35-25) and then frame-projected through a 165mm focal length tube lens (Zeiss) onto the CMOS camera (Photometrics Prime 95B). The excitation arm consists of a 473 nm laser (OptoEngine, BL-II-473/1~100mW) coupled to a collimator (Thorlabs, F230FC-A). A 50 mm lens was then used to focus the beam through an dichroic mirror (Semrock, FF495-Di02-25×36) onto the back focal plane of the objective. The path was manually aligned to ensure uniform illumination of the three fiber bundles. The laser power was adjusted so that 0.25 mW was emitted, measured at the end of each fiber tip, in order to minimize photobleaching over multiple extended recording sessions.

#### Optical Characterization

##### Point spread function estimation

Five microliters of stock solution 0.2-μm-diameter fluorescent beads (Invitrogen, FluoSpheres Sulfate Microspheres, 0.2 μm, yellow-green fluorescent (505/515)) was mixed in a solution of 1.5% aqueous agarose (Sigma). This mixture was then poured onto a glass-bottomed petri dish and was allowed to solidify at room temperature. A single imaging fiber was held above the dish and manually lowered using a linear translation stage (Thoralbs, MT1A) and used to acquire images of the illuminated beads at 5 μm intervals (Figure S7A). Intensity along X, Y, and Z dimensions were quantified using ImageJ.

##### Linear resolution

A single fiber was used focused above a standardized 1951 USAF Resolution Test Target (Thorlabs). After acquiring, the smallest pair of three lines was identified and linear intensity quantified in ImageJ (Figure S7B).

##### SNR comparison

To ensure that the use of multiple fiber cores did not diminish the SNR of recorded Ca^2+^ transients *in vivo*, a single mouse was injected with AAV1-CaMKIIa-GCaMP6f in ACC and a 0.6mm GRIN lens (Inscopix) was implanted 0.2mm above the injection site, as described above. After 3 weeks, the mouse was head fixed and imaged using our fiber bundle system at 34 Hz over 5 minutes. The mouse was then imaged through the same objective and imaged with a CMOS camera placed in the widefield path of a Bruker microscope, illuminated with the same laser power and wavelength (0.25 mW, 473nm) again for five minutes. Individual sources were extracted and registered across the fiber bundle and the standard widefield recordings (Figure S7C, top; see **Data Acquisition & Post-processing)**. Recordings of the same cells were overlaid and qualitatively appeared to be similar in quality (Figure S7C, bottom left). Peak-to-noise ratio was calculated for each cell for each imaging method and found to be quantitatively similar (R = 0.61) (Figure S7C, bottom right).

### Data Acquisition & Post-processing

#### Image acquisition

Prior to each session, mice were headfixed and each GRIN lens was carefully cleaned with 70% ethanol. Custom 3D printed adaptors were first affixed to each lens, and then fibers were manually adjusted vertically while acquiring images until cells were maximally focused. Because both the fibers and lenses had the same diameter of 0.6mm, the design of the adaptors ensured that minimal horizontal adjustments were required to attain the same field-of-view across multiple recordings. Images were collected during behavior at 34 Hz (exposure time 17 ms) using the Photometrics data acquisition software, Programmable Virtual Camera Access Method (PVCAM) at a resolution of 920 pixels/mm.

##### Paired Optogenetic Inhibition during training and GCaMP imaging during retrieval

During training, bilateral optogenetic inhibition of AM was paired with GCaMP6f imaging through a unilaterally-placed GRIN lens in the ACC (Figure S9A) during training sessions, using our previously described fiber bundle microscope. Inhibition was achieved with a 473 nm laser (OptoEngine) via a mono fiber patch cord with a minimum power of 15 mW, and was targeted through the bilateral implanted cannulas in AM for the entire time the mouse was in the cue zone (8 seconds). Imaging was performed during the pre-exposure session, R6 and R27 sessions. Imaging laser power was maintained at 0.25 mW to minimize distal inhibition of GtACR. To observe the online effects of AM inhibition on ACC neural activity, an additional recording session was performed after the first pre-exposure session, where imaging was paired simultaneously with inhibition. The mouse was head-fixed in the behavior box in the dark, and eight separate 8-second optogenetic laser pulses occurring 30 seconds apart were delivered while imaging occurred concurrently through the GRIN lens (Figure S9B).

##### Source extraction

Each field of view was cropped and motion-corrected using NoRMCorre (Pnevmatikakis and Giovannucci, 2017). A combination of custom scripts and existing packages for 1-photon excitation source extraction (CNMF-E, Zhou et al., 2018) were used to identify and isolate individual cell ROIs and dF/F signals from all three regions separately. Cell identification was verified by manually validating every extracted source. Cell registration across sessions was performed with a combination of custom scripts and existing packages (Cell Reg, Sheintuch et al., 2017).

##### Calculation of single cell dF/F and transient identification

For each cell detected via automated source extraction, a normalized ΔF/F was calculated and individual Ca^2+^ transients were identified as previously described (Rajasethupathy et al., 2015). Briefly, ΔF/F was defined as: (F - F_baseline_)/F_baseline_, where F is the raw output (“*C_raw”*) from the CNMF-E algorithm, and where F_baseline_ is the baseline fluorescence, calculated as the mean of the fluorescence values for a given cell, continuously acquired over a 20 s sliding window to account for slow time-scale changes observed in the fluorescence across the recording session. For all analysis, this dF/F was then normalized by z-scoring the entire time series across a session. To identify statistically significant transients, we first calculated an estimate of the noise for each cell using a custom MATLAB script, with a previously described method. In essence, we identified the limiting noise cutoff level for a given cell using time periods that are unlikely to contain neural events, and then defined a transient as significant if it reached above at least 3σ of this estimated noise level. A custom MATLAB script using the function *findpeaks* was used to identify any remaining obvious transients not identified by this method (typically when multiple transients occurred in rapid succession). Additional specifications required transients to persist above this noise level for at least 300 ms (roughly twice the duration of the half-life decay time of GCaMP6f). The transient duration was defined as the first and last frames where the dF/F reached 3σ. The value of dF/F was set to zero outside the duration of every identified transient to minimize effects of residual background fluorescence.

##### Correlations

To calculate cell-to-cell correlations (either inter- or intra-regional), we calculated Pearson’s R for each pair of time series. To minimize unwanted correlations that may occur from long-tailed Ca^2+^ transients, we only included the dF/F values from a cell during the rise time and one second after the peak for each transient. We defined the transient rise as the frames from when the dF/F value first rises above the noise threshold (see <u>Calculation of single cell dF/F and transient identification</u>) until the frame when the dF/F reaches a maximal value within the identified transient time window. The rest of the time series was set to zero. When correlations were calculated for a given context (i.e., high reward correlations), the time series for each cell was concatenated to only include the frames that were captured during the cue zone for that context. For calculating correlations of HR-tuned cells, the cue zones of both contexts were concatenated before calculating Pearson’s R.

A cell pair was defined as highly-correlated if R>0.3. Correlation network graphs were displayed using the MATLAB function *graph* with a line connecting correlated cells, and the layout set to force-directed. Best match correlation was defined as the maximum R value achieved for a given cell in one region when correlated to cells in another region or with other cells in that same region. To measure the distribution of interconnectedness within a region during a recording session, we calculated the number of intra-regional correlated cell pairs achieved by each cell (network degree/cell) and then normalized by dividing the total number of cells in that recording. When selecting cells that were highly correlated with cells in a different region (AM – ACC or HPC – ACC), we selected cells from the top 5% R-values from each intra-regional cell pair on that day. Chance distributions for Figure 6K were calculated by randomly selecting the same number of cells as there are red cells from the network graph and re-calculating the number of highly correlated cell pairs per cell 1000 times. Significance corresponds to a proportion of the distribution outside of the 95% percentile of this chance level. Distributions include values from all cells from all animals within a given region and/or experimental condition.

##### Single cell contextual tuning

To calculate the tuning of an individual cell to the high reward over the low reward context for a given session we calculated the difference in number of significant transients occurring during the cue zone in concatenated high reward trials and low reward trials, each normalized by the total time spent in the cue zone. Thus the units are events/second and a positive value corresponds to increased average event rate in the high over the low reward context. For each cell, we also calculated a “shuffled” tuning, where the time between transients across the entire session was randomly shuffled and tuning was recalculated for a given cell a total of 1000 times. A cell was considered high reward-tuned if the magnitude of its un-shuffled tuning was at least 1 standard deviation above its shuffled distribution of tunings.

### Statistical Analyses

Sample sizes were selected based on expected variance and effect sizes from the existing literature, and no statistical methods were used to determine sample size a priori. Prior to experiments being performed, mice were randomly assigned to experimental or control groups. The investigator was blinded to all behavioral studies. Data analyses for calcium imaging (in vitro and in vivo datasets) were automated using MATLAB scripts. Statistical tests were performed in MATLAB 2017a, 2021b, or Graphpad Prism. Animals were excluded from experiments using the following criteria: For all behavioral experiments, mice were excluded if by T5, they had a greater rate of anticipatory licking in the aversive condition compared to reward conditions (indicating they had not properly learned the task).

### Data & Code availability

Custom MATLAB scripts developed and used in this study will be available as a Github repository on our lab website.

---

**Figure S1.**
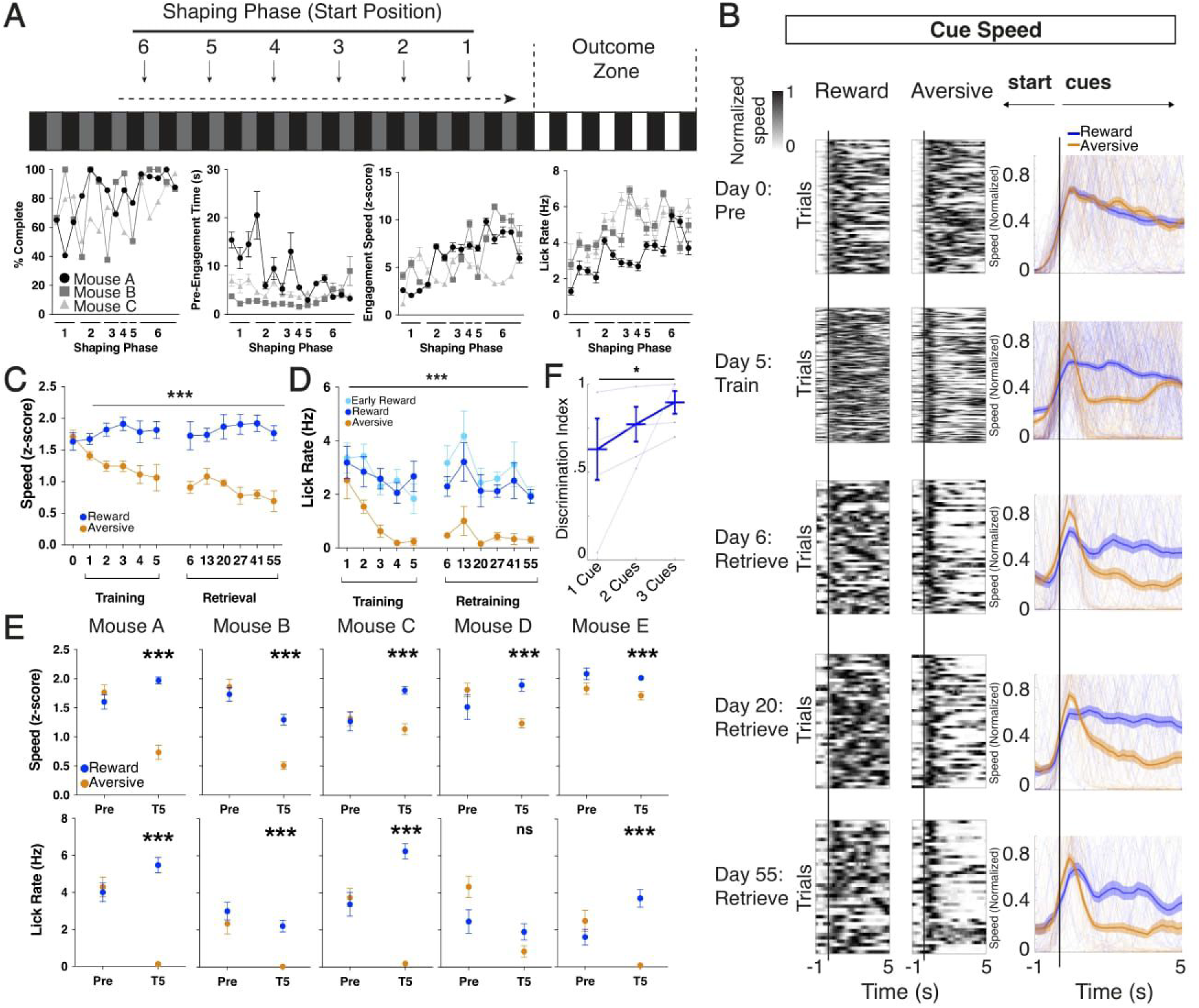
Individual mouse performance and speed contextual discrimination, related to Figure 1. A. Virtual reality linear track design showing starting position of the mouse for each phase of the shaping protocol. Plots for three representative mice during shaping showing percentage of trials complete, time (s) spent prior to engagement, speed (z-score) during engagement, and anticipatory lick rate (Hz) upon entry to outcome zone. B. Raster of individual trials and plot of trial averages. Speed prior to and during cue zone. Data in heatmaps are normalized to peak speed in the trial. N=5 mice. C. Average speed in cue zone in each context during preexposure, training, and weekly retrieval sessions. N=5 mice, ***p<0.001 for speed, Two way ANOVA with repeated measures, data are mean± s.e.m. D. Anticipatory lick rate (Hz) 2 seconds prior to outcome zone in reward trials at the start of each training session (light blue) is comparable to that of the remainder of the trials in that session (dark blue). Also shown is lick rate on retraining sessions between reward (dark blue) and aversive (orange), following each retrieval session, which have comparable behavioral performance to training sessions. N=5 mice, ***p<0.001, Two way ANOVA with repeated measures, data are mean± s.e.m. E. Speed (z-score) and lick rate (Hz) in cue zone between reward and aversive, shown individually for each mouse for preexposure and training day 5. ***p<0.001, unpaired t-test, data are mean± s.e.m. F. Quantification of discrimination between reward and aversive anticipatory lick rates per mouse when 1, 2 or 3 cue modalities are presented after training day 5. N=4 mice. *p<0.05 between 1 cue and 3 cues, Dunn’s multiple comparison, data are mean± s.e.m.

**Figure S2.**
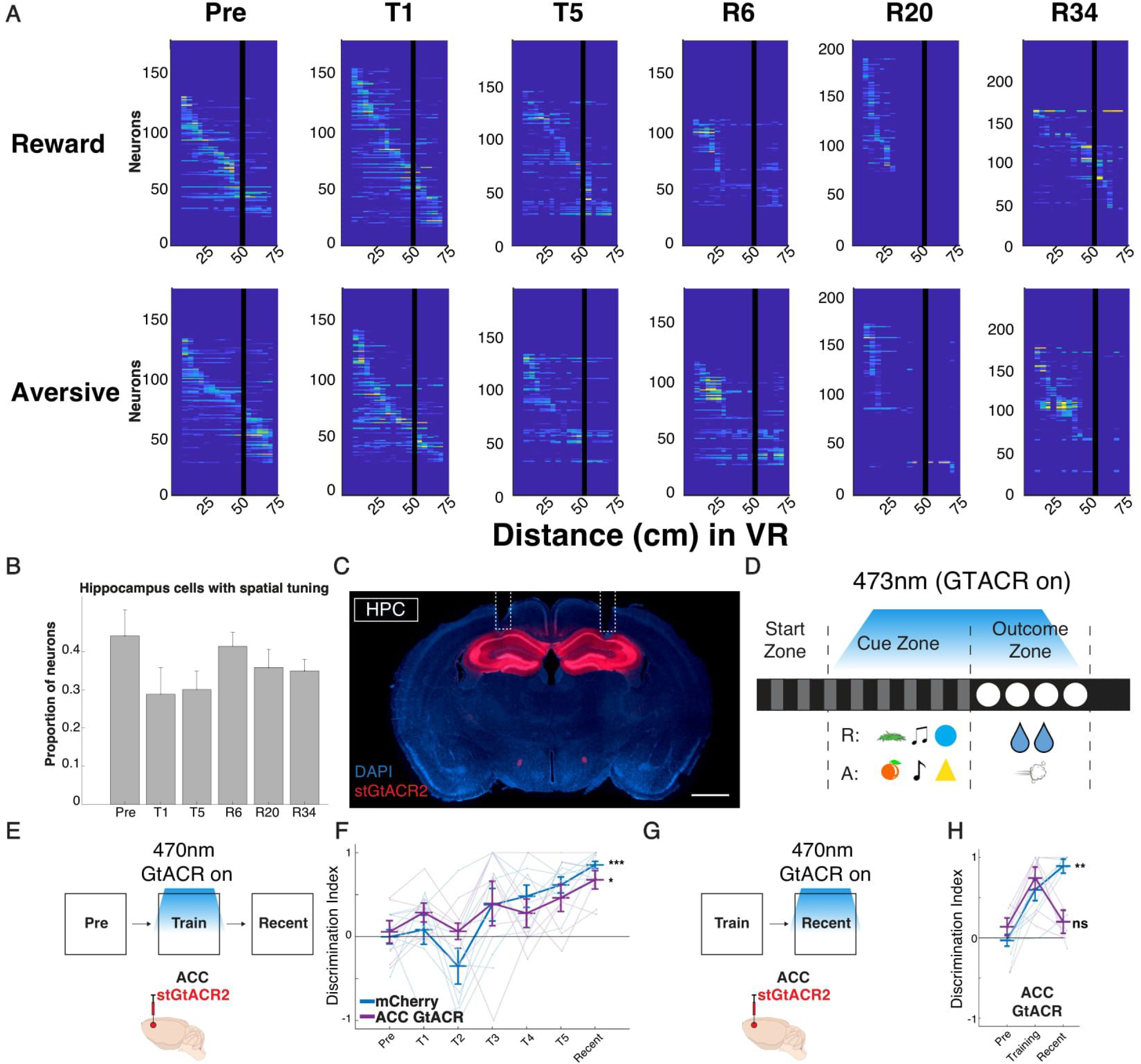
Hippocampus place cell activity and histology for ACC and HPC optogenetic cohorts, related to Figures 1 and 2. A. Recorded dF/F from hippocampal cells (see Methods, In Vivo Multi-Region Calcium Imaging) binned by location on the virtual track B. Proportion of cells exhibiting spatially-tuned responses, as defined by mean dF/F on the track when binned by spatial location exceeding one standard deviation above mean dF/F during the inter-trial interval (n=3 mice). C. Coronal sections from animals virally injected with AAV1-CamKii-stGtACR2 bilaterally in HPC. White tracts denote location of fiber optic implants. DAPI is shown in blue and stGtACR2 in red. Scale: 1mm. D. Schematic showing light for optogenetic inhibition was delivered during cue and outcome periods of the trial. E. Schematic of experimental design: stGtACR2-based optogenetic inhibition of ACC during training (T1-T5), followed by a test of recent memory. F. Quantification of discrimination between reward and aversive lick rates per mouse cohort on preexposure, training, and recent retrieval sessions. ***p<0.001 between pre-exposure and recent for mCherry vs p=0.0212 for ACC GtACR, one-way repeated measures ANOVA with Tukey’s multiple comparison test. Individual data points shown, with mean± s.e.m. G. Schematic of experimental design: GtACR-based optogenetic inhibition of ACC during recent memory in animals that were trained without optogenetic inhibition. H. Quantification of discrimination between reward and aversive lick rates per mouse cohort on preexposure, training, and recent retrieval sessions. **p<0.01 for between pre-exposure and recent for mCherry vs p=0.921O for ACC GtACR, one-way repeated measures ANOVA with post-hoc Tukey’s multiple comparison test. Individual data points shown, with mean± s.e.m.

**Figure S3.**
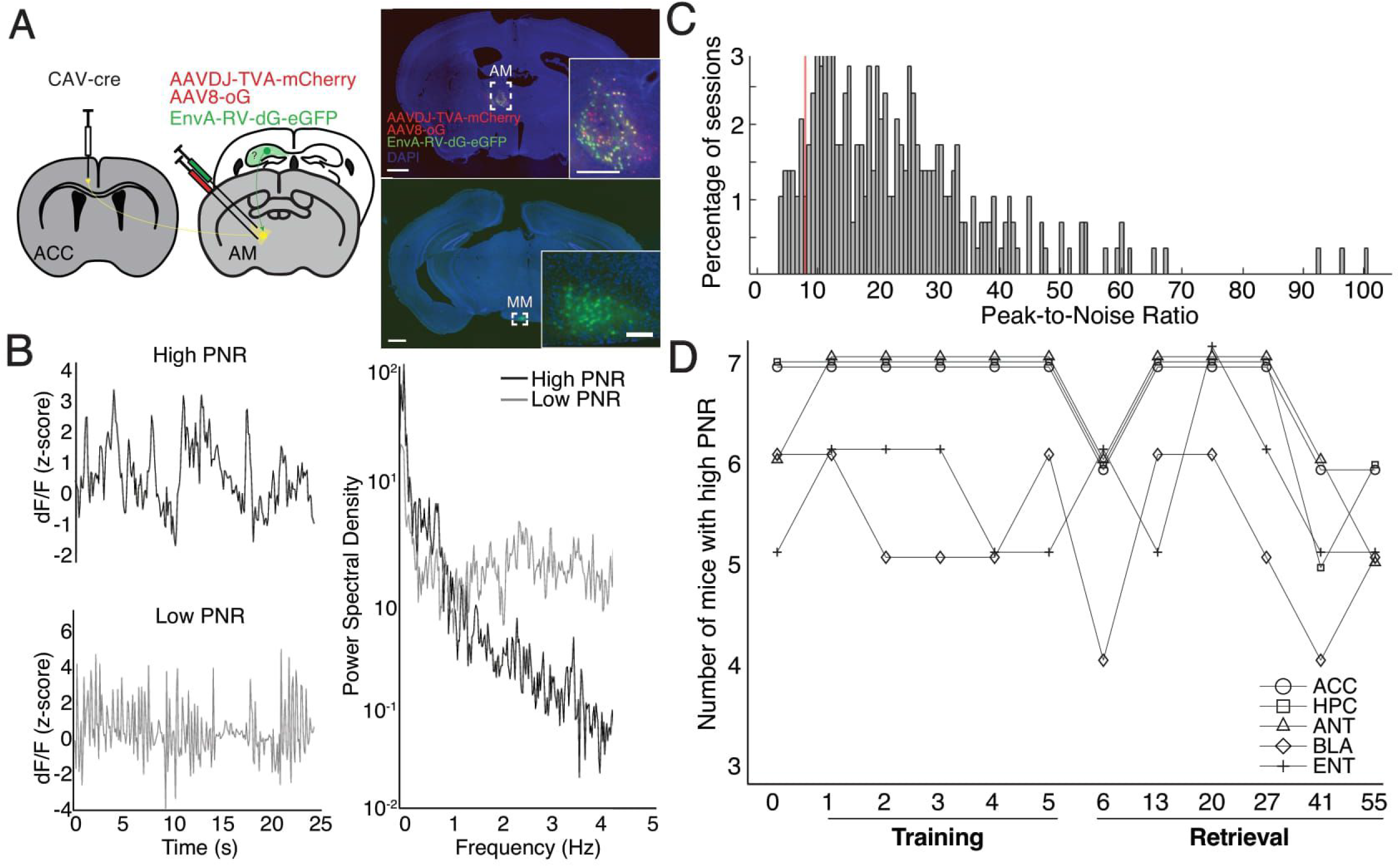
TRIO tracing and signal-to-noise of photometry data, related to Figure 3. A. Strategy for trans-synaptic tracing of input to AM-ACC neurons. CAV-cre was injected into ACC along with AAVs expressing ere-dependent TVA-mCherry (TC)/rabies glycoprotein (G) into AM, followed by glycoprotein deleted and GFP-expressing rabies viruses (RVdG). Expression in mammillary bodies (MM) is shown. Scale: 1 mm or 200um for inserts. B. Left: Examples of a photometry recording with a high (black) or low (gray) peak-to-noise ratio (PNR). Right: The high PNR recording consists of high amplitude, low-frequency fluctuations lasting several seconds, while the low PNR recording consists of sharp, high-frequency fluctuations. C. Distribution of PNR for all recording sessions, and empirically selected noise threshold shown in red. D. Total number of recordings (i.e., number of mice) included from each region and for each session in the final analyses.

**Figure S4.**
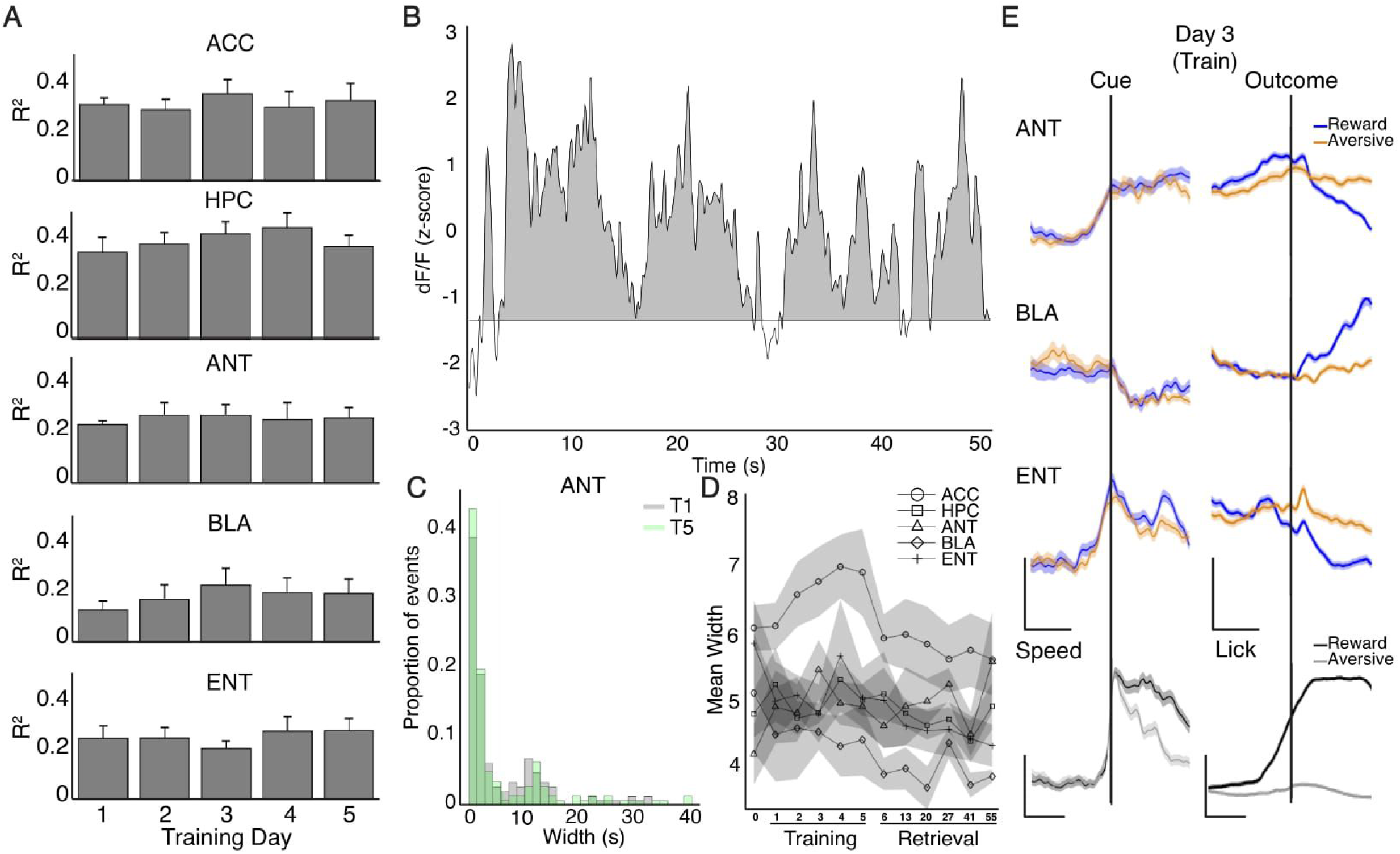
Event rates and regression data for photometry signals, related to Figures 3 and 4. A. Mean full-model R2 calculated for photometry recordings from each region during training (see methods). N=7 mice. Data are mean± s.e.m. B. Example photometry signal demonstrating large, slow calcium events lasting tens of seconds. C. Distribution of all photometry signal event durations from ANT on the first (T1) and last (TS) day of training. N=7 mice. D. Mean calcium event width across mice for each region and recording session. N=7 mice. Data are mean ± s.e.m. E. Photometry signals (dF/F, z-score) and behavioral variables speed (z-score) or lick rate (Hz) aligned to cue zone or outcome zone entry, respectively for day 3 of training (T3). N=7 mice, dark lines and shaded regions represent mean± s.e.m. Photometry scale: x/y: 2s/1z.

**Figure S5.**
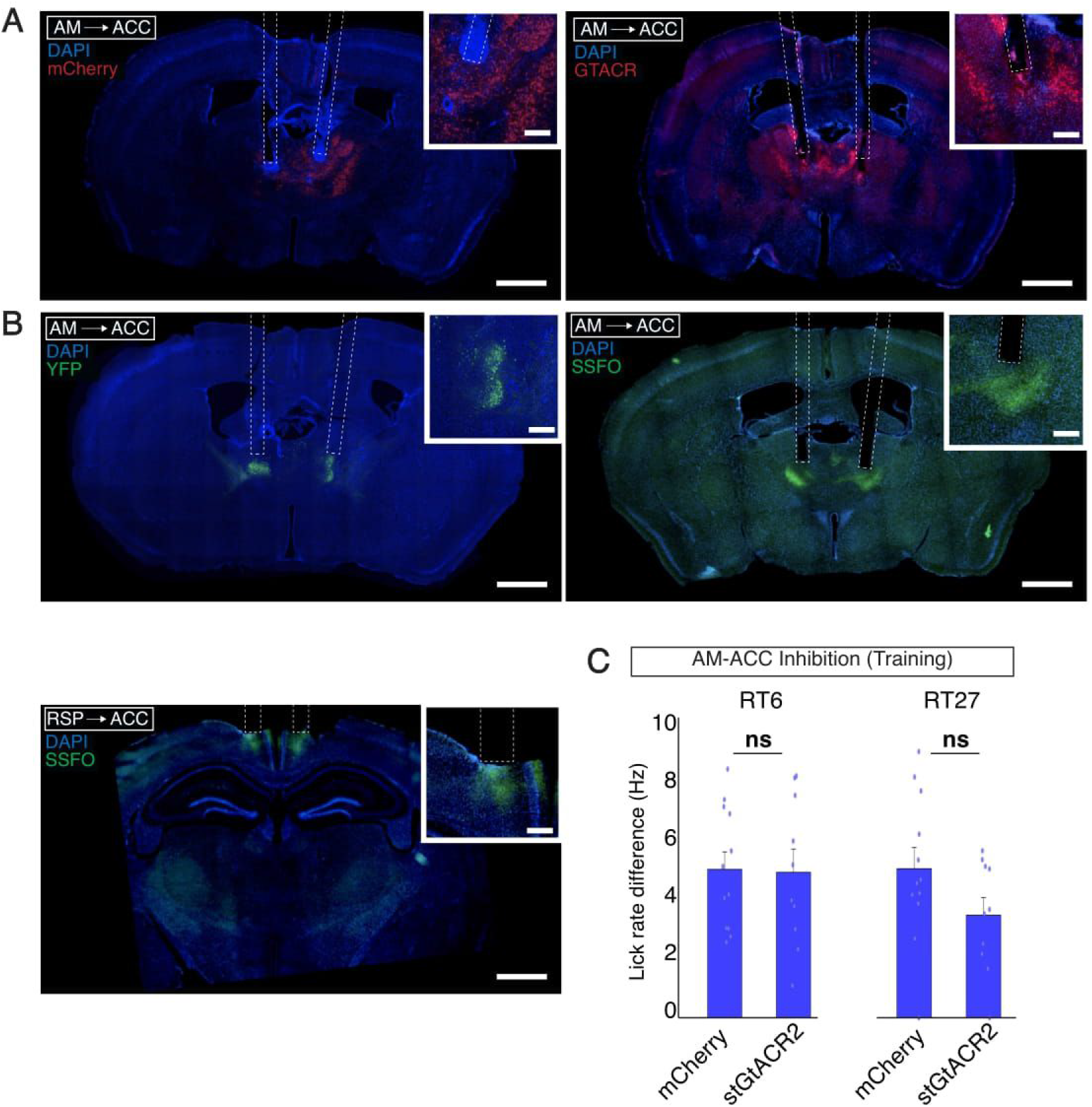
Histology and retraining behavior for optogenetic cohorts, related to Figure 5. A. Coronal sections from animals virally injected with rgAAV-hSyn-Cre bilaterally in ACC and either AAV1-SIO-stGtACR2-FusionRed or AAV 1-DIO-mCherry bilaterally in AM. White tracts denote location of fiber optic implants. Insert is a zoomed in image of viral expression. DAPI is shown in blue and mCherry or stGtACR2 is shown in red. Scale: 1mm or 200um for insert. B. Coronal sections from animals virally injected with rgAAV-hSyn-Cre bilaterally in ACC and either Ef1a-DIO-SSFO-EYFP or hSyn1-DIO-eYFP bilaterally in AM (above) or Ef1a-DIO-SSFO-EYFP bilaterally in retrosplenial cortex (RSP; below). White tracts denote location of fiber optic implants. Insert is a zoomed in image of viral expression. DAPI is shown in blue and YFP or SSFO is shown in yellow. Scale: 1mm or 200um for insert. C. Behavioral data from AM-to-ACC inhibition during training experiment, quantifying performance on each retraining session following retrieval for all mice in Figures 5A-D, shown as lick rate differences (Hz) in reward vs aversive contexts. N=12 mice for mCherry vs 10 mice for stGtACR2, all p>0.05, unpaired t-test. Individual data points shown, as well as mean ± s.e.m.

**Figure S6.**
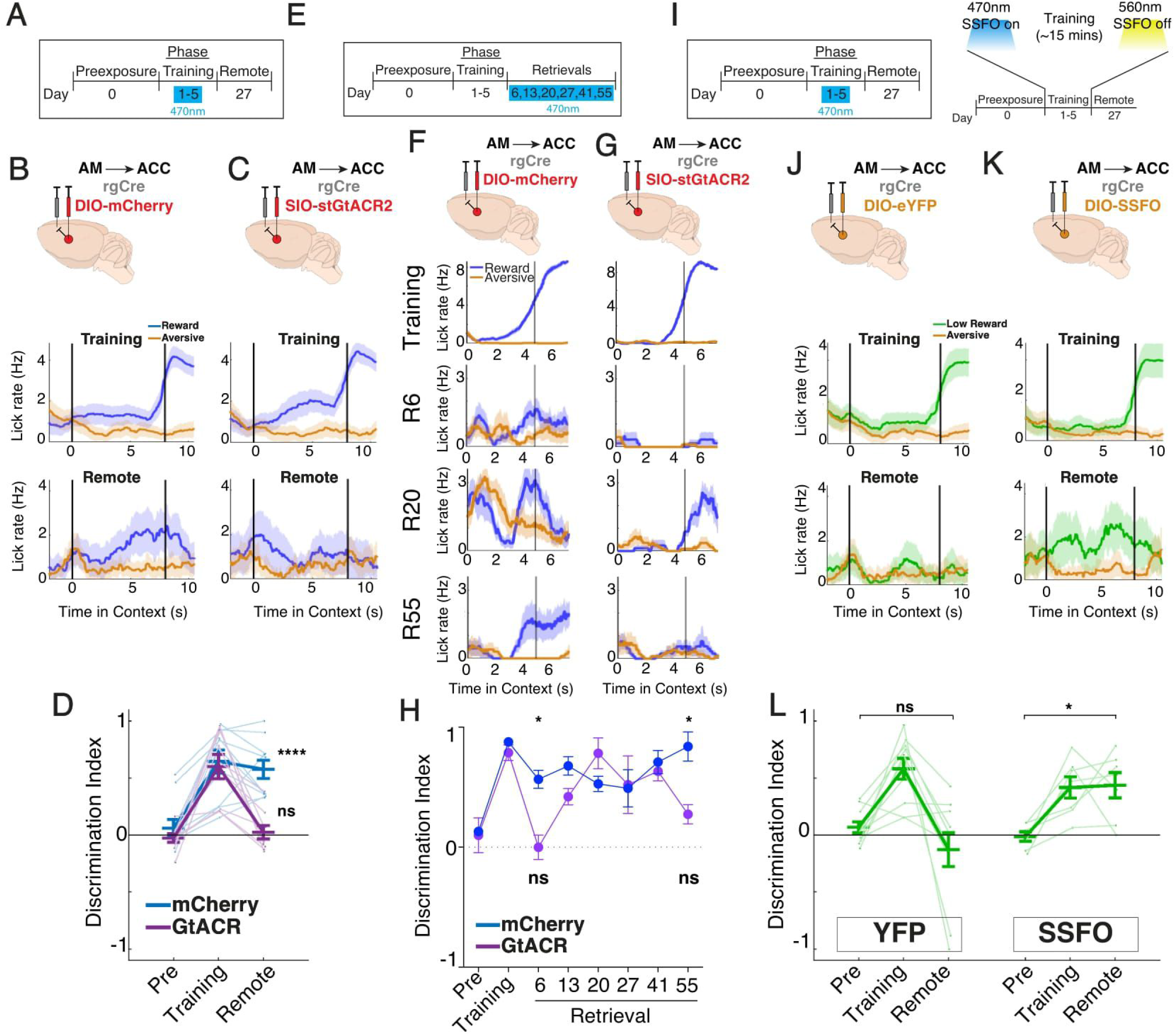
Optogenetic perturbations of AM->ACC activity without recent retrieval and retraining, related to Figure 5. A. Schematic of experimental design: stGtACR2-based optogenetic inhibition during training (T1-T5), followed by a test of remote (R27) memory. Light was delivered during cue and outcome periods of the trial. B. C. Injection strategy for targeting anteromedial thalamus (AM) projections to ACC (AM->ACC) in mCherry control and stGtACR2 opsin cohorts. Raw lick traces in each context on training day 5 and remote retrieval sessions. D. Quantification of discrimination between reward and aversive lick rates per mouse on preexposure, training day 5, and remote retrieval sessions; mCherry (AM->ACC no opsin control, N=10), GtACR (AM->ACC with opsin, N=9). ****p<0.0001 for mCherry between preexposure and remote, one-way repeated measures ANOVA with post-hoc Tukey’s multiple comparison test. Individual data points shown, with mean ± s.e.m. E. Schematic of experimental design: stGtACR2-based optogenetic inhibition during retrieval probes (R6, R13, R20, R27, R41, and R55). Light was delivered during cue and outcome periods of the trial. F, G. Injection strategy for targeting anteromedial thalamus (AM) projections to ACC (AM->ACC) in mCherry control and stGtACR2 opsin cohorts. Raw lick traces in each context on training day 5, R6, R20, and R55 retrieval probe sessions. H. Quantification of discrimination between reward and aversive lick rates per mouse on preexposure, training day 5, R6, R20, and R55 retrieval sessions; mCherry (AM->ACC no opsin control, N=7), GtACR (AM->ACC with opsin, N=3). *p<0.05 for mCherry between Preexposure and R6 or Preexposure and R55 sessions. p>0.05 for GtACR, one-way repeated measures ANOVA with post-hoc Tukey’s multiple comparison test. Individual data points shown, with mean± s.e.m. I. Schematic of experimental design: SSFO-based enhancement of neural excitability during training (T1-T5), followed by testing of remote memory. 470nm light was delivered at the start of each training session to activate the SSFO and then 560nm light delivered at the end to deactivate the SSFO. J,K. Injection strategy for targeting AM projections to ACC in YFP control and SSFO opsin cohorts. Raw lick traces in each context on training day 5 and remote retrieval sessions. L. Quantification of discrimination between low reward (LR) and aversive (A) contexts on preexposure, training day 5, and remote sessions for each cohort; YFP (no opsin control, N=10); SSFO (AM->ACC SSFO excitation, N=6). *p<0.05 for YFP and SSFO cohorts between preexposure and remote, one-way repeated measure ANOVAs with post-hoc Tukey’s multiple comparison tests. Individual data points shown, with mean± s.e.m.

**Figure S7.**
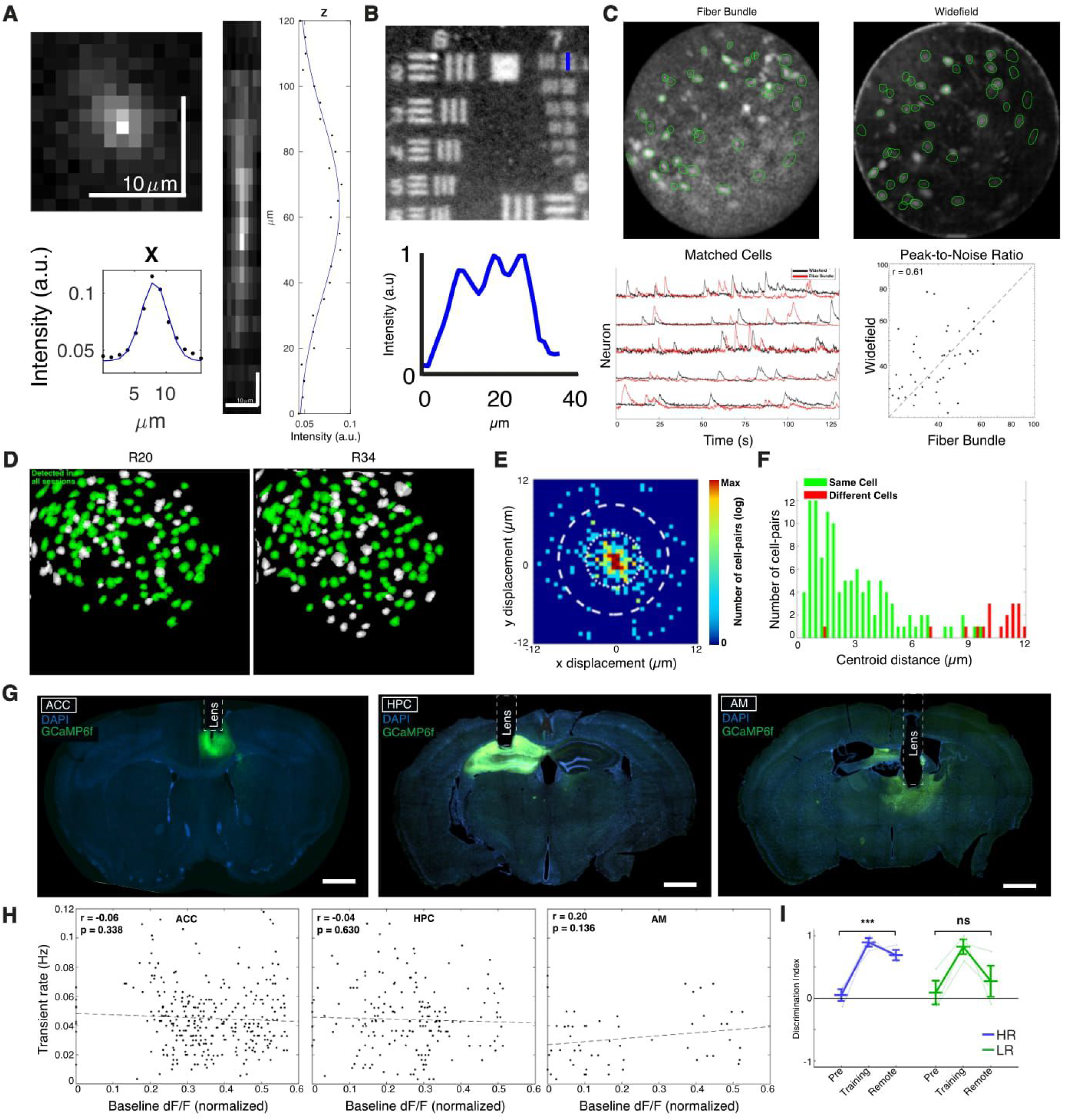
Fiber bundle microscope characterization, in-vivo imaging histology and behavior, related to Figure 6. A. Projections and quantification of a three-dimensional stack of observed fluorescence from a sub-resolution fluorescent microbead acquired using the fiber bundle microscope. B. Left: Fiber bundle-acquired image of a USAF imaging target. Right: Quantification of intensity along the group with the finest resolved spatial frequency. C. Top: Snapshot of an in vivo recording of ACC with the fiber bundle microscope (left) and a standard widefield microscope (right), with registered cells circled in green. Bottom left: Overlayed dF/F time series from 5 example cells recorded with the two microscopes. Bottom right: Comparison of estimated Peak-to-Noise ratio (from CNMF-E) of each cell plotted against itself from the two methods. D. Spatial footprints on R20 (left) and R34 (right); footprints identified in both session are labeled by green. E. Post-alignment offset of identified matched spatial footprints. F. Centroid distance histograms for matched (green) and non-matched (red) pairs of spatial footprints. G. Brain histology from a representative mouse in the 3-way multi-region imaging cohort showing GCaMP6f expression and GRIN lens implantation site in ACC, HPC, and AM. Scale: 1 mm. H. Spontaneous event rate of GCaMP6f-expressing neurons as a function of baseline fluorescence intensity in each region. N=3 mice. Dotted line: best fit line. Lack of correlation suggests that calcium buffering (i.e., increasing levels of baseline GCaMP) in these regions does not affect physiological event rates. I. Discrimination between high reward (HR) or low reward (LR) and aversive (A) contexts in the cohort of imaged mice. N=3 mice. *p<0.05 for HR between preexposure and remote, one-way repeated measure ANOVA with post-hoc Tukey’s multiple comparison test. Individual data points shown, with mean± s.e.m.

**Figure S8.**
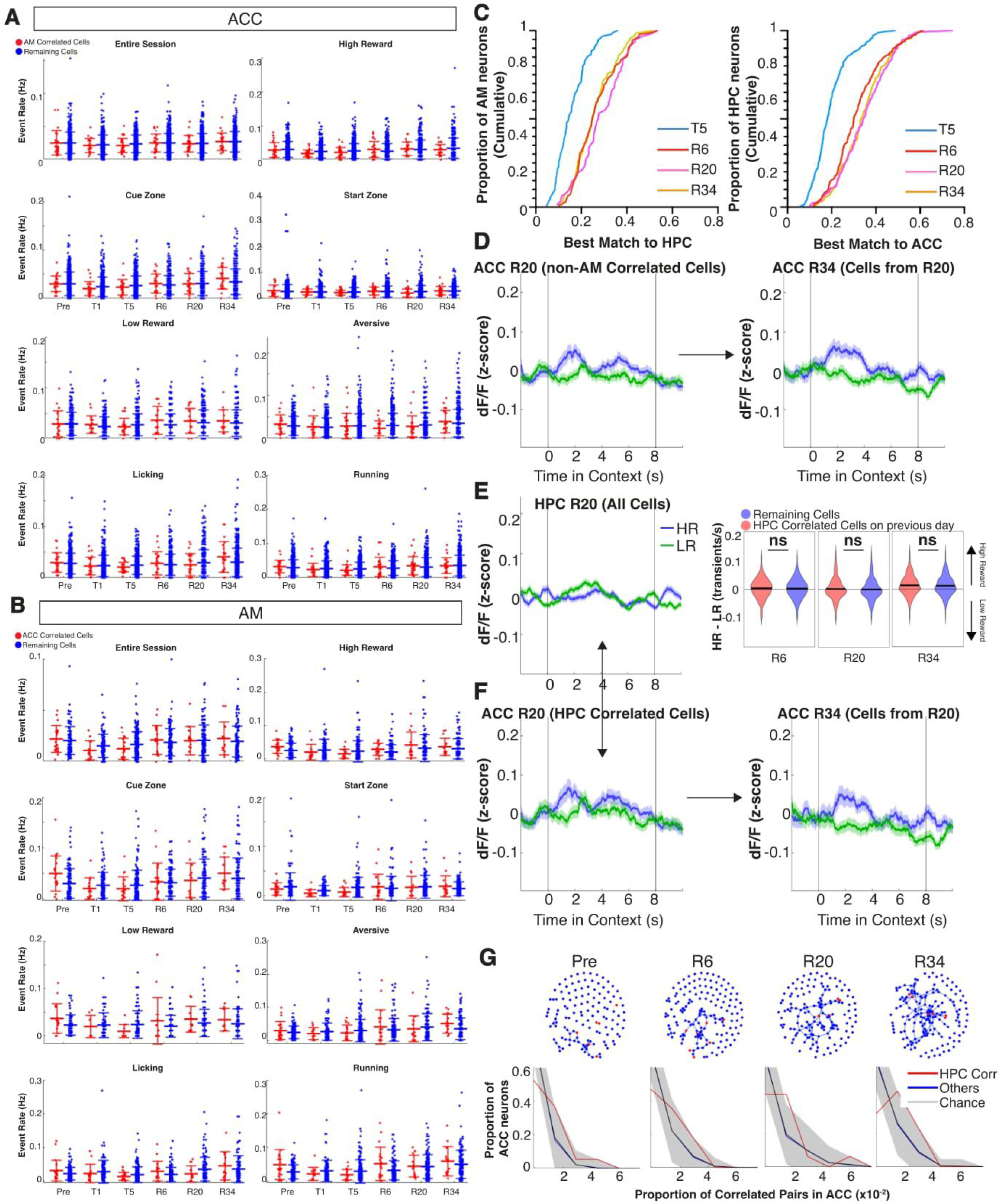
Cross-region event rates, functional correlations, and synchronous activity, related to Figure 6. A. Event rate (transients per second) of ACC cells which are highly correlated to AM cells (red, top 10%), and all remaining ACC cells (blue) during context-specific cue zones in the task. For “running” or “licking", transients are counted any time speed or licking, respectively, are greater than zero. N=all cells from 5 mice. Data are mean± s.e.m. B. Same as S6A, but for the AM cells identified in the top 10% highly correlated AM-to-ACC pairs, compared to all remaining AM cells. N=all cells from 5 mice. Data are mean± s.e.m. C. Cumulative distribution of the best match correlation of HPC cells paired with AM cells across days (left) and ACC cells paired with HPC cells (right). N=5 mice. p>Cl.05, two-sample KS test between R6 and R20. D. Left: Cue-zone-aligned mean z-scored dF of all ACC cells on R20 not correlated to an AM cell (n=242 cells). Right: mean response of these same tracked ACC cells on R34. E. Cue-zone-aligned mean z-scored dF of all HPC cells on R20 (n=518 cells). Bottom left: Mean z-scored response of ACC cells which form highly correlated pairs with at least one HPC cell (n=171, cells across 3 animals. Bottom right: mean response of these same tracked ACC cells on R34. F. Distribution of tuning for ACC cells correlated to at least one HPC cell on the previous recording session compared to the tuning of all remaining cells. The difference was not significant on R34, R20 or R6. G. Top: Example from one animal of undirected network graphs of ACC cell correlations (Lines are Pairwise Pearson>Cl.3) in the reward context across days with ACC cells most correlated with HPC cells (top 5%) shown in red (see methods). Bottom: Distribution of proportion of pairwise correlations in ACC for red cells and blue cells, compared to chance shown in black (see methods). N=5 mice, Shaded area represents 95% confidence interval around chance distribution.

**Figure S9.**
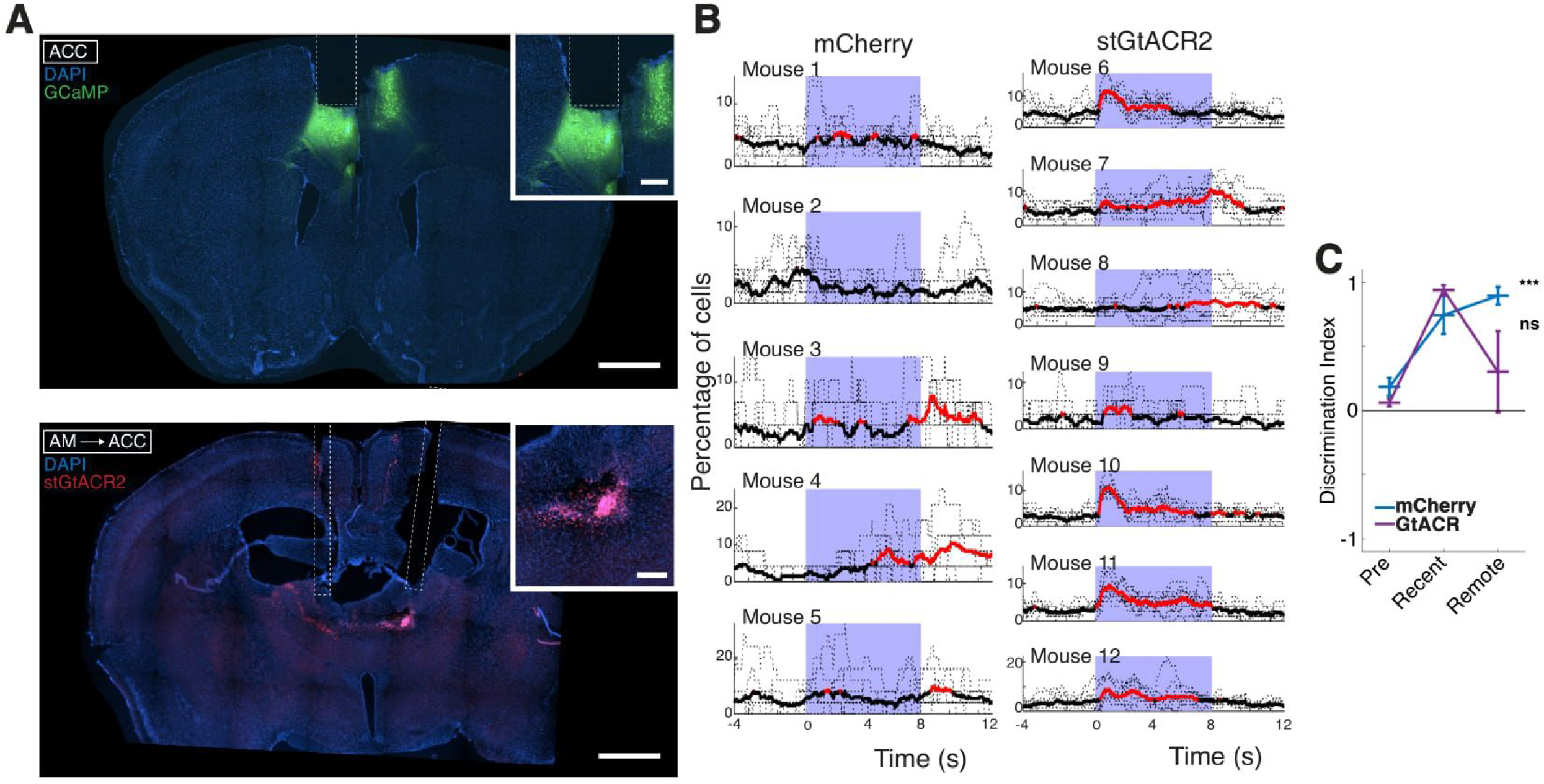
Histology and characterization of paired inhibition and imaging cohort, related to Figure 7. A. Brain histology from a representative mouse undergoing paired inhibition (AM-to-ACC) and imaging (ACC), from Fig. 4J-M. Coronal sections show GCaMP6f expression and GRIN lens implantation site in ACC, and stGIACR2-FusionRed expression and fiber optic tracts in AM. Insert is a zoomed in image of viral expression. DAPI is shown in blue, GCaMP6f is shown in green, and stGtACR2 is shown in red. Scale: 1 mm or 200um for insert. B. Percentage of cells having a significant transient event at every frame, averaged across mice, for mCherry and stGtACR2 cohorts. Red denotes time points with synchronous firing greater than 1 s.d. above the mean across 8 pulses. Purple highlights duration of laser on (470nm) for stGtACR2 activation. A. Discrimination between reward and aversive contexts for mice undergoing paired inhibition and imaging. ***p<0.001 for mCherry between preexposure and remote, one-way repeated measure ANOVA with post-hoc Tukey’s multiple comparison test. Individual data points shown, with mean ± s.e.m.

